# A bistable autoregulatory module in the developing embryo commits cells to binary fates

**DOI:** 10.1101/2022.10.31.514335

**Authors:** Jiaxi Zhao, Mindy Liu Perkins, Matthew Norstad, Hernan G. Garcia

## Abstract

Positive autoregulation has been repeatedly proposed as a mechanism for cells to adopt binary fates during embryonic development through bistability. However, without quantitatively determining their parameters, it is unclear whether the plethora of positive autoregulatory modules found within developmental gene regulatory networks are actually bistable. Here, we combine *in vivo* live imaging with mathematical modeling to dissect the binary cell fate dynamics of the fruit fly pair-rule gene *fushi tarazu* (*ftz*), which is regulated by two known enhancers: the early (non-autoregulating) element and the autoregulatory element. Live imaging of transcription and protein concentration in the blastoderm revealed that binary Ftz cell states are achieved as *ftz* expression rapidly transitions from being dictated by the early element to the autoregulatory element. Moreover, we discovered that Ftz concentration alone is insufficient to activate the autoregulatory element, and that this element only becomes responsive to Ftz at a prescribed developmental time. Based on these observations, we developed a dynamical systems model, and quantitated its kinetic parameters directly from experimental measurements. Our model demonstrated that the *ftz* autoregulatory module is indeed bistable and that the early element transiently establishes the content of the binary cell fate decision to which the autoregulatory module then commits. Further analysis *in silico* revealed that the autoregulatory element locks the Ftz expression fate quickly, within 35 min of exposure to the transient signal of the early element. Overall, our work confirms the widely held hypothesis that autoregulation can establish developmental fates through bistability and, most importantly, provides a framework for the quantitative dissection of cellular decision-making based on systems dynamics models and real-time measurements of transcriptional and protein dynamics.

## 1 Introduction

One of the central questions in developmental biology concerns how cells precisely and irreversibly adopt distinct cellular fates. It has been argued that cells assume their unique gene expression profiles through a sequence of decisions among branching paths (***Zernicka-Goetz et al., 2009***; ***Soldatov et al., 2019***), most famously encapsulated by C. H. Waddington’s “epigenetic landscape” of peaks and valleys delineating the possible trajectories that a cell can follow (***Waddington, 1957***). Genetic networks that lock a cell into one of these trajectories may be thought of as “memory modules” that guide cells through valleys in the landscape to their ultimate fates. In the simplest case, where a decision is made between two alternative developmental fates, the memory module is binary and is often referred to as a switch. The state of the switch is set by the action of transient upstream regulatory signals.

Several genetic motifs, such as autoactivation and mutual repression, have been identified that are capable of maintaining binary cell fates (***Alon, 2007***; ***Peter and Davidson, 2015***). However, the mere presence of a motif is insufficient to guarantee that a network can remember its expression state once upstream regulators have degraded. The ability to lock onto high or low expression levels results from bistability (***Box 1***), a systems-level property that depends upon the quantitative details of the kinetics of the involved chemical reactions (***Ferrell, 2002***; ***Angeli et al., 2004***; ***Graham et al., 2010***).

Despite the widespread invocation of bistability to explain the stable and irreversible determination of cellular fates (***Peter and Davidson, 2015***), relatively little quantitative data exist to confirm bistability in gene expression modules within developing embryos. Previous studies in cell culture and fixed embryos have provided evidence for the existence of bistability in hematopoietic differentiation (***Laslo et al., 2006***; ***Kueh et al., 2013***), the Shh network (***Lai et al., 2004***), the vertebrate hindbrain (***Bouchoucha et al., 2013***), between the BMP and FGF morphogens (***Srinivasan et al., 2014***), and within the Notch-Delta signaling system (***Sprinzak et al., 2010***). Quantitative evidence for multistability in fruit fly embryos has also been derived from fitting the parameters of high-dimensional network models to measurements in fixed tissue (***von Dassow et al., 2000***; ***Jaeger et al., 2004***; ***Lopes et al., 2008***; ***Manu et al., 2009***; ***Papatsenko and Levine, 2011***; ***Verd et al., 2017***). While these models are capable of reproducing the observed phenomenology, there is often no guarantee that the optimal set of inferred parameter values reflects actual biophysical quantities (***Gutenkunst et al., 2007***; ***Cotterell and Sharpe, 2010***; ***Villaverde et al., 2015***; ***Wieland et al., 2021***). Thus, it is important to verify that the conclusions drawn from computational modeling and *in vitro* experiments apply to developmental systems *in vivo* in the context of models that quantitatively capture the molecular interactions that underlie the process of cellular decision making. To the best of our knowledge, evidence for the bistability of a genetic module based on these molecular interactions in an intact multicellular organism has not yet been demonstrated.

The early development of the fruit fly *Drosophila melanogaster* is an ideal model system for studying binary cell fate decision making. Specifically, the mapping of the regulation of *fushi tarazu* (*ftz*), one of the *Drosophila* pair-rule genes that forms seven discrete stripes at the cellular blastoderm stage prior to gastrulation (2.5 - 3.5 hours after fertilization; ***Nüsslein-Volhard and Wieschaus (1980***); ***Hafen et al. (1984***); ***Wakimoto et al. (1984***); ***Weiner et al. (1984***); ***Hiromi et al. (1985***)), has suggested that this gene is capable of autoactivation (***Hiromi and Gehring, 1987***). These studies showed that two main enhancers—the early, or zebra, element and the autoregulatory element— dictate Ftz protein expression during early embryogenesis (***Figure 1***A; ***Hiromi and Gehring (1987***)). The early element responds to upstream transcription factors such as the gap genes to establish the initial expression pattern of seven stripes (***Dearolf et al., 1989***). This element is functionally distinct from the autoregulatory element, which contains multiple Ftz binding sites that allow Ftz to activate its own expression (***Pick et al., 1990***; ***Schier and Gehring, 1992, 1993***). This autoactivation network motif is theoretically capable of exhibiting bistability and has therefore been hypothesized to act as a binary memory module (***Alon, 2007***; ***Xiong and Ferrell, 2007***).

**Figure 1.**
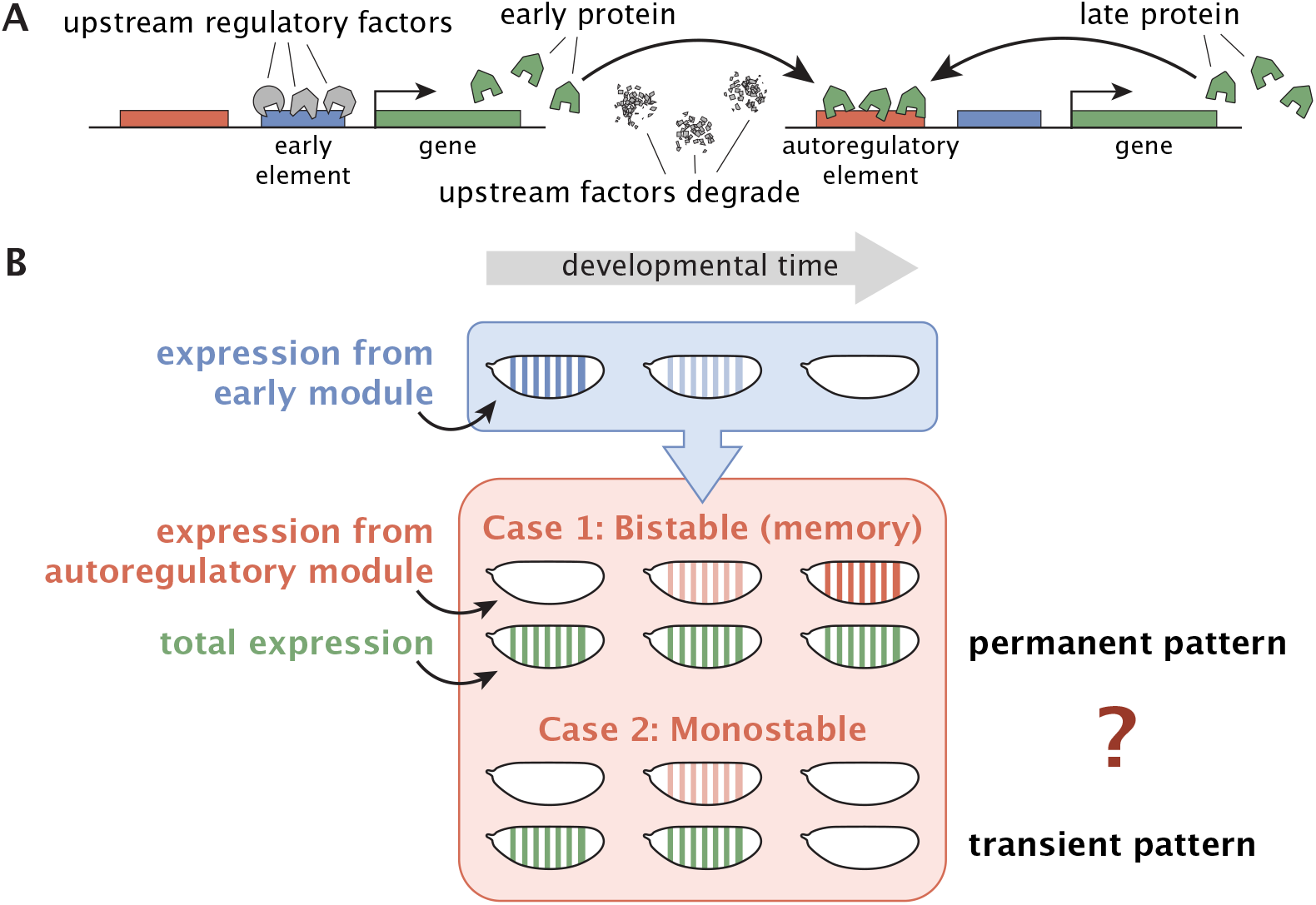
The bistability of the autoregulatory module can determine cell fate. (**A**) The autoregulatory architecture consists of an early element to which upstream factors bind to transiently upregulate gene expression of an activator, and an autoregulatory element to which the activator binds to, under certain circumstances, promote its own expression even once the upstream factors have degraded. (**B**) The total activator concentration (green) is a sum of the protein produced by the early (blue) and autoregulatory (red) modules. If the autoregulatory module is bistable, it possesses binary memory that permits transiently high concentrations of early protein to be locked into permanent high expression levels (high cell fate). If, in contrast, the autoregulatory module is monostable, then it may transiently boost protein levels from the early module, but over time all cells will ultimately revert to the same low fate.

Whether a cell possesses a memory module determines whether observed states of gene expression are transient in the absence of continued external signaling, or whether these states can be locked into permanent cell fates that can be maintained without further intervention. Specifically, if the autoregulatory module is bistable, then high *ftz* expression driven by the transient presence of upstream factors is stabilized by the autoregulatory element into a permanent cell fate (***Figure 1***B, case 1), even once those factors degrade (or until further regulatory mechanisms intervene). If, instead, the autoregulatory element is monostable, then the observed separation of Ftz concentration into high and low levels persists only as long as upstream factors are present to regulate expression. In their absence, Ftz expression would revert to a single fate for all cells (***Figure 1***B, case 2). It is important to note, however, that in this case, the transiently high or low trajectory of Ftz concentration could still be instructive for regulating downstream genes.

Here, we characterize the *ftz* autoregulatory module *in vivo* through quantitative real-time measurements in living fruit fly embryos. We focus on the anterior boundary of stripe 4, the only Ftz stripe that has been shown to be driven exclusively by the early and autoregulatory elements and not by other enhancers in the gene’s vicinity (***Schroeder et al., 2011***; ***Graham et al., 2021***). We observe that Ftz expression separates into high and low levels at the blastoderm stage during the 20 min prior to gastrulation, concurrent with a transition in regulatory control from the early to the autoregulatory element. We discover that autoregulation is triggered at a specific time point in development—presumably through the action of “timer genes” (***Clark and Akam, 2016***; ***Clark and Peel, 2018***; ***Clark et al., 2022***)—rather than through a readout of Ftz concentration alone. Based on these observations, we develop a dynamical systems model and quantitate its parameters from simultaneous real-time measurements of *ftz* transcription and Ftz protein dynamics in single cells of living embryos. Our model predicts binary Ftz expression levels at gastrulation with high accuracy and demonstrates that, indeed, the *ftz* autoregulatory module is bistable. We conclude that the *ftz* autoregulatory element acts as a memory module to commit cells to binary fates that are otherwise transiently defined by the early element, thereby validating a long-standing hypothesis in developmental and systems biology. Simulations further make it possible to quantitatively define a developmental commitment window, which shows that the autoregulatory module requires about half an hour to establish a memory of the transient signal from the early module. Thus, our work provides a framework for the dissection of other regulatory modules in the gene regulatory networks that dictate development based on this interplay between dynamical systems models and real-time experiments.

## 2 Results

### 2.1 Binary cell states of Ftz expression are established in early development

To understand the role of positive autoregulation in deciding Ftz expression levels, we first sought to visualize and track Ftz protein dynamics over time. We used CRISPR-mediated recombination (***Gratz et al., 2015***) to fuse a LlamaTag, a fluorescent probe that reports on the fast protein dynamics that characterize early embryonic development, to the C-terminus of the endogenous Ftz protein (***Figure 2***A; ***Bothma et al. (2018***)). An examination of the fluorescently labeled Ftz protein in the early embryo shows that, around 15 min before gastrulation, Ftz protein is expressed in a seven-stripe pattern with clear, smooth boundaries (***Figure 2***B, left). This expression pattern refines over the following 15 min into sharp stripe boundaries by the start of gastrulation (***Figure 2***B, right). The result shows that cells express either high or low levels of Ftz protein, as pictured in ***Figure 2***C for the anterior boundary of stripe 4, consistent with results from previous studies (***Schroeder et al., 2011***; ***Clark, 2017***; ***Bothma et al., 2018***).

**Figure 2.**
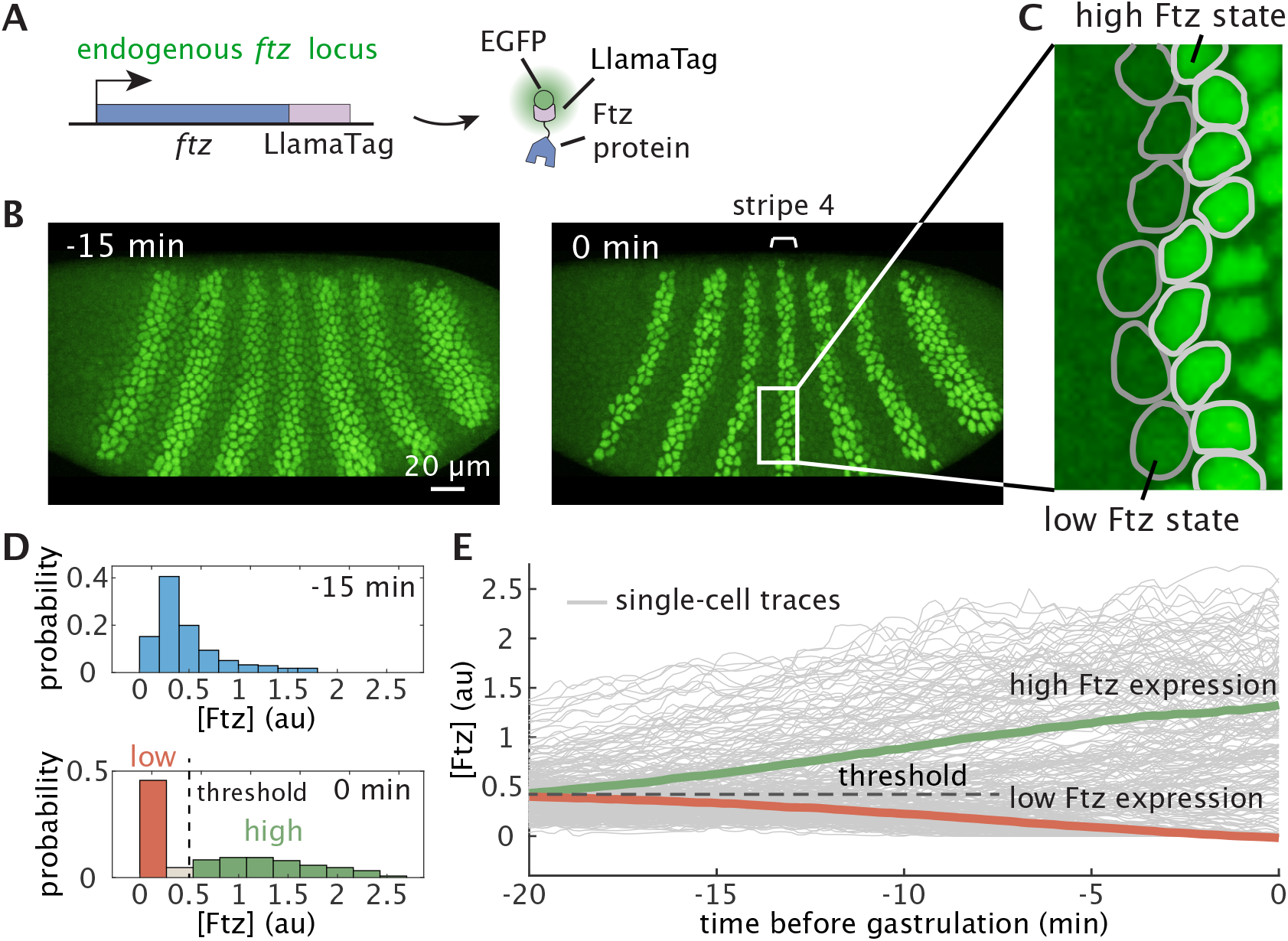
Binary cell states are rapidly established prior to gastrulation. (**A**) A fusion of endogenous Ftz to a LlamaTag makes it possible to visualize the highly dynamic Ftz protein pattern in the early fly embryo. Once Ftz protein is translated in the cytoplasm, LlamaTag binds to maternally deposited EGFP and is transported into the nucleus, increasing nuclear fluorescence to produce a direct readout of Ftz protein concentration. (**B**) Snapshots from a movie capturing Ftz protein concentration dynamics. The anterior side of the embryo is oriented towards the left, and the time is given relative to gastrulation. (**C**) Ftz expression along the anterior boundary of stripe 4 shows a discrete transition between distinct cell states. (**D**) Histograms of single-nucleus fluorescence values at different developmental time points show that a single threshold can be used to classify cells into “high-Ftz” and “low-Ftz” cell states prior to gastrulation. (**E**) Single-cell trajectories of nuclei at the anterior boundaries of Ftz stripe 4. Green and red lines are averages for nuclei determined to ultimately have “high-Ftz” and “low-Ftz” levels at gastrulation, respectively, as defined in (D).

Our live imaging measurements allowed us to quantitatively examine the dynamics with which binary cell states are established by calculating the Ftz protein distribution in individual nuclei at different time points in development (***Figure 2***D). Our analysis revealed that the expression level is initially unimodal across all cells (***Figure 2***D, top and E) and then evolves into a bimodal distribution within 15 min (***Figure 2***D, bottom and E). Consistent with our qualitative observations in ***Figure 2***B and C, cells at these later times can be quantitatively classified into distinct “high-Ftz” and “low-Ftz” cell states using a single threshold (***Figure 2***D, bottom and E), which indicates that binary cell states are already established prior to the onset of gastrulation. Moreover, though the two cell states are clearly distinguishable from each other, we observed that there is significant cell-to-cell variability within both states. Specifically, the single-nucleus Ftz protein distribution for high-Ftz levels spans more than a two-fold range (***Figure 2***D, bottom and E).

Previous studies have established that autoregulation plays a key role during *ftz* expression: a lack of the *ftz* autoregulatory element or mutated Ftz binding sites within the element result in the loss of Ftz expression at later developmental stages (***Hiromi and Gehring, 1987***; ***Schier and Gehring, 1992***). However, it is unclear at what developmental time *ftz* autoregulation is initiated in response to the Ftz expression stemming from the early element. Specifically, is the autoregulatory element active before stripes of Ftz expression emerge, or is autoregulation invoked after the early element has already established this pattern? To distinguish between these two scenarios, we decoupled the transcriptional dynamics driven by the early and autoregulatory elements by creating two separate reporter constructs, each containing only the early or autoregulatory elements followed by MS2 stem-loops that enable the direct visualization of transcriptional dynamics (***Figure 3***A; ***Garcia et al. (2013***); ***Lucas et al. (2013***)).

**Figure 3.**
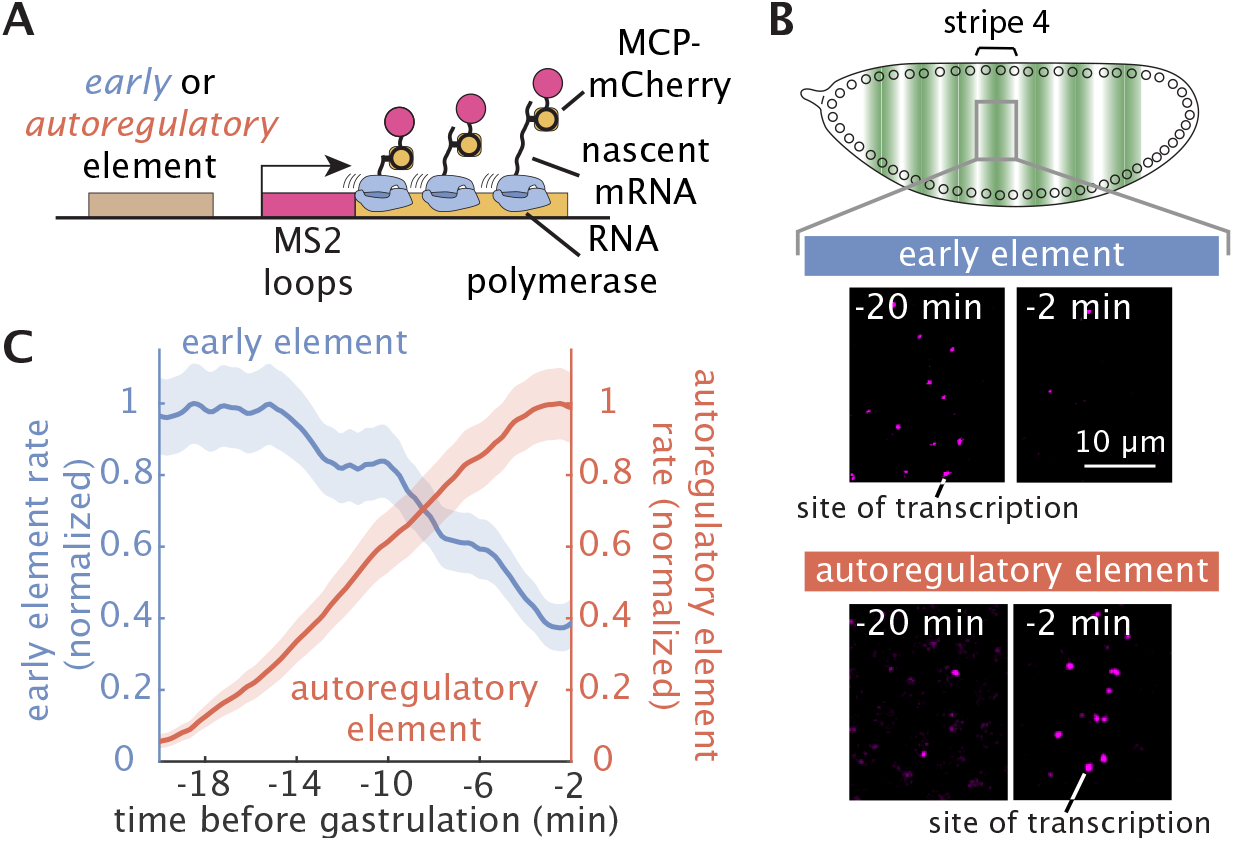
A sequential transition in Ftz regulation from the early to the autoregulatory element occurs during the establishment of discrete cell states. (**A**) Imaging transcriptional dynamics of the early and autoregulatory elements using the MS2 system. Maternally deposited MS2 coat protein (MCP) fused to mCherry binds to MS2 stem-loops in the nascent RNA of the reporter construct. (**B**) Snapshots of sites of nascent transcript formation labeled by MS2 from reporters of the early and autoregulatory elements at different time points reveal that transcription from the early element is reduced significantly as gastrulation approaches and that transcription driven by the autoregulatory element increases shortly before gastrulation. (**C**) Quantification of the transcriptional activity reported by the MS2 fluorescence from the early (*N* = 3 embryos) and autoregulatory (*N* = 7 embryos) elements as a function of time confirms that, within 20 min, *ftz* gene expression transitions from originating mainly from the early element to being dominated by the autoregulatory element. MS2 traces are smoothened using a moving average of 5 min. Error bars shown indicate standard errors over multiple embryos.

#### Box 1.

**Bistability**

A simplified dynamical systems model of the *ftz* autoregulatory element that ignores the dynamics of mRNA production describes the rate of change in Ftz concentration over time as

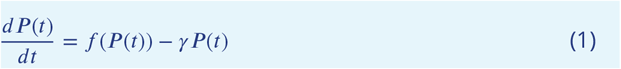

where *P* is Ftz concentration, *f* (*P*) is the gene regulatory (or input-output) function that describes how input Ftz concentration controls the rate of Ftz production, and *γ* is the Ftz degradation rate. Since Ftz promotes its own production, *f* (*P*) increases with *P*.

In the long term, Ftz concentrations will tend toward stable *steady states*, or attractors, for which (by definition) the change in concentration over time goes to 0 (i.e., 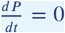). Attractors are stable, meaning that if Ftz concentration is perturbed slightly away from the attractor, it will eventually return to the attractor. However, it is also possible to have unstable steady states for which, after a small perturbation, Ftz concentration will evolve away from the steady state. For the system in ***Eq. 1***, a steady state *P*^*^ will solve

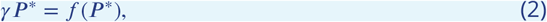

meaning that all steady states can be found graphically as the intersections between a line of slope *γ* and the function *f*. In our case, the gene regulatory function *f* is a sigmoidal function (red curves in Figure B1), such that there are between 1 and 3 intersections of the total degradation rate *γP* with the Ftz production rate *f* (*P*). Then the autoregulatory module is either monostable, meaning it possesses one attractor, or bistable, meaning it possesses two attractors and one unstable steady state between these attractors. Since the number of intersections depends on the shape of *f* and the slope *γ*, the exact parameter values are crucial for determining the possible behaviors that can be exhibited by the autoregulatory module.

**Figure B1.**
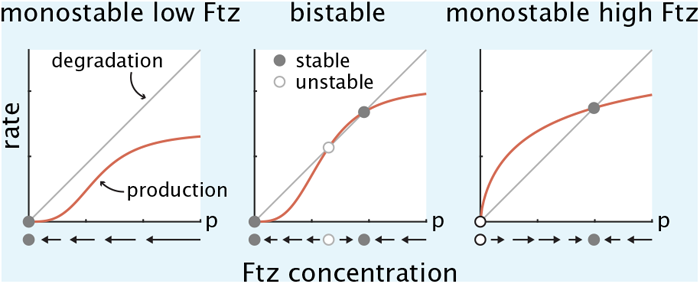
Above, steady states are identified through the intersection of the production rate *f* (*P*) with the total degradation rate *γP* where *p* is the Ftz concentration. Below, vector fields show the direction Ftz concentration will evolve over time. An equivalent graphical test also exists for 2D dynamical systems models where mRNA is modeled explicitly; see ***Section S1.3***.

We will also consider controlled systems, i.e., dynamical systems that have an additional regulatory input. In this case, this input corresponds to the Ftz concentration produced by the early element, *P*_*early*_, such that the dynamics of Ftz production by the autoregulatory element are given by

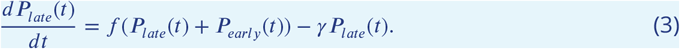

Since the early element shuts down over time (that is, *P*_*early*_(*∞*) = 0), the steady states *P*_*late*_ in ***Eq. 3*** are the same as those for *P* in ***Eq. 1***. However, the transient dynamics of *P*_*early*_ shape the trajectory *P*_*late*_ will take and determine which steady state Ftz will ultimately reach.

We observed that the early element already drives a relatively constant gene expression level around 20 min prior to gastrulation (***Figure 3***B and C). Then, at 15 min before gastrulation, its transcriptional activity decreases significantly, resulting in a 60% reduction within the next 20 min of development (***Figure 3***B and C). Conversely, autoregulation is initiated 20 min prior to gastrulation, with its activity increasing until gastrulation starts (***Figure 3***C). This transition between the early and autoregulatory elements occurs while binary cell states are being established (***Figure 2***). Since autoregulation becomes the dominant driver of Ftz expression after this transition, it is likely that autoregulation plays a key role in Ftz-mediated decision-making prior to gastrulation.

### 2.2 *ftz* autoactivation is triggered at a specific developmental time

The tight transition between the early and autoregulatory elements that occurs within 20 min (***Figure 3***) could be indicative of autoactivation being initiated by the increase in Ftz concentration driven by the early element. However, autoactivation could also be triggered by upstream factors at a specific developmental time, regardless of the Ftz level at that time point.

To distinguish between these two scenarios, we measured the gene regulatory function—the input-output function describing how the input Ftz concentration dictates the output rate of *ftz* transcription—of the *ftz* autoregulatory element at distinct developmental times. If autoactivation is solely initiated by Ftz produced by the early element, the regulatory function should remain constant throughout development. On the other hand, if autoregulation is triggered through an independent mechanism—such as activation mediated by other transcription factors at a given developmental time—then the *ftz* autoregulatory element should be unresponsive to input Ftz protein before this developmental time point.

We measured the gene regulatory function of the autoregulatory element by constructing an experimental system that allows for the simultaneous monitoring of Ftz concentration and the corresponding autoregulatory activity. We used the tagged endogenous Ftz protein as the input and introduced a transgenic reporter with MS2 loops under the control of the *ftz* autoregulatory element as the output (***Figure 4***A). Live imaging of the anterior boundary of Ftz stripe 4 (***Figure 4***B) showed that, initially, around 25 min prior to gastrulation, both Ftz expression and the autoregulatory response were relatively low, and later increased as development progressed. Just before gastrulation, the Ftz protein pattern refined into a discrete boundary, with the *ftz* autoregulatory response clearly following the stripe boundary (***Figure 4***B).

**Figure 4.**
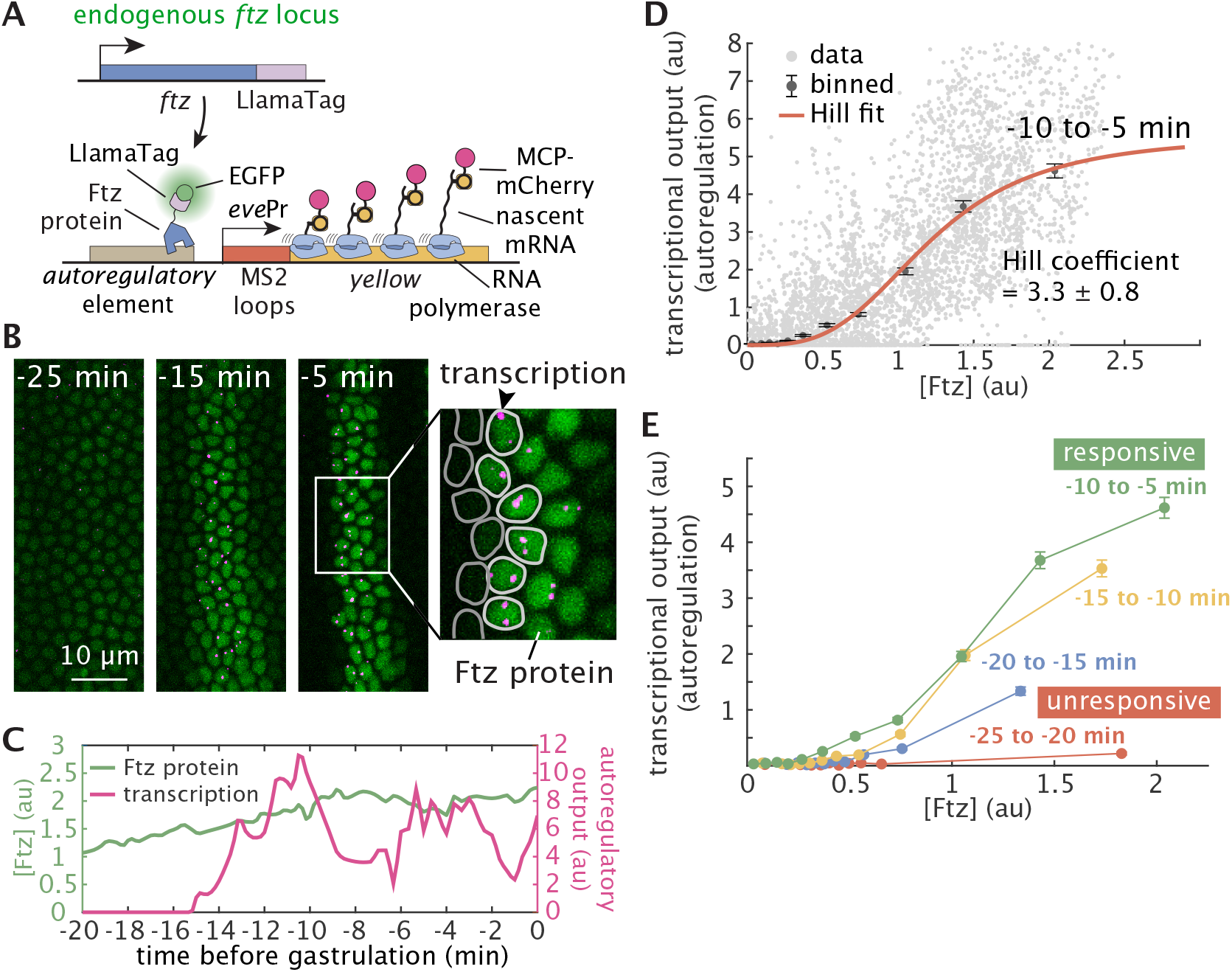
Two color live-imaging reveals that *ftz* autoregulation is initiated at a specific developmental time. (**A**) Two-color tagging permits *in vivo* simultaneous visualization of input Ftz protein concentration using a LlamaTag and output autoregulatory transcriptional dynamics using a reporter carrying the MS2 system. (**B**) Representative frames from live-imaging data. Green and magenta channels correspond to Ftz concentration and the transcriptional output from the *ftz* autoregulatory element, respectively. Dark gray and light gray outlines correspond to cells expressing high or low levels of Ftz protein, respectively. (**C**) Illustrative single-cell trace of Ftz protein and autoregulatory activity. Green and magenta lines correspond to the Ftz protein and transcriptional activity of the autoregulatory element, respectively. Both protein and MS2 traces are smoothened using a moving average of 1 min. (**D**) Experimentally measured gene regulatory function of the *ftz* autoregulatory element between -10 min to -5 min relative to gastrulation. Grey points correspond to simultaneous measurements of Ftz and MS2 fluorescence at individual time points from single-cell traces at the anterior boundary of stripe 4 (*N* = 211 nuclei, example traces shown in ***Figure S1***). These points were grouped into quantiles, and a Hill function (red line) was fit to the quantile means. (**E**) The autoregulatory input-output function evolves over time, as the *ftz* autoregulatory element transitions from an unresponsive to a responsive state within 15 min, indicating that *Ftz* autoregulation is initiated through a developmental time-based mechanism. Error bars shown indicate standard errors. All data are from *N* = 7 embryos.

To calculate the regulatory function, we restricted our analysis to the cells at the anterior boundary of stripe 4 as a means to minimize the influence of other position-dependent transcription factors that might also contribute to *ftz* autoactivation (***Schier and Gehring, 1993***). We first extracted two rows (high and low) of boundary cells. Then, we separated the input Ftz concentration and output transcription from the autoregulatory element in the data corresponding to each individual cell (***Figure 4***C; ***Figure S1***) into ten quantiles and fit a Hill function to the quantile averages to get the gene regulatory function of *ftz* autoregulation within a defined temporal range (for example, -10 min to -5 min for ***Figure 4***D). Our analysis revealed a sharp regulatory relationship between Ftz protein and autoregulatory response, with a Hill coefficient of 3.3±0.8 (***Figure 4***D). Such Hill coefficients are comparable to those estimated in the context of autoactivation in vertebrate hindbrain development (***Bouchoucha et al., 2013***) as well as those observed in simpler regulatory motifs that do not feature feedback (***Gregor et al., 2007***).

We repeated the process described above at multiple developmental times to analyze how the regulatory function for *ftz* autoactivation evolves over time. The results, shown in ***Figure 4***E, revealed that the regulatory function is clearly distinct at different time points. Specifically, initially around -25 min to -20 min, the *ftz* autoregulatory element is effectively unresponsive to input Ftz protein (***Figure 4***E, red line). However, the element progressively transitions to a fully responsive state within 15 min (***Figure 4***E, green line). The observed temporal evolution of the autoregulatory element’s responsiveness to Ftz protein is a clear indication that the autoregulatory element is not always primed to respond to input Ftz protein and that, instead, its expression is triggered at a specific developmental time, presumably by upstream transcription factors.

### 2.3 Mathematical modeling quantitatively predicts Ftz concentrations

As we argued in ***Figure 1***B, the fact that Ftz can exhibit a high state at one point in time does not necessarily imply that this high state will persist in the absence of upstream regulation. Determining whether the autoregulatory module is bistable and hence possesses developmental memory requires turning the schematic shown in ***Figure 1***A into an explicit mathematical model with empirically determined parameter values. To that end, we first developed a dynamical systems model for the full Ftz regulatory system (including both the early and autoregulatory elements) to verify whether we could accurately recapitulate experimental results *in silico*.

Our measurements revealed that the autoregulatory element is unresponsive to Ftz concentration until about *t*_*on*_ = −20 min (***Figure 4***E). Once the element becomes responsive, we can describe the *ftz* autoregulatory module using the dynamical systems model given by

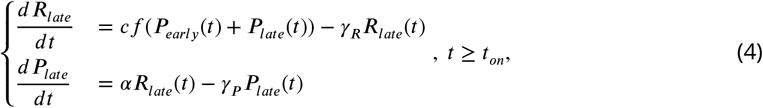

where *R*_*late*_(*t*) and *P*_*late*_(*t*) are the mRNA and protein concentrations produced by the autoregulatory module, *γ*_*R*_, *γ*_*P*_ are the decay rates of mRNA and protein, respectively, and *α* is the translation rate. *f* (*P*) is the gene regulatory function measured for the transgene and shown in ***Figure 4***D that describes the output rate of mRNA production as a function of the input Ftz concentration. Since both the early and autoregulatory elements drive Ftz expression, the gene regulatory function depends on *P*_*total*_(*t*) which is the sum of the protein contributions from the early (*P*_*early*_(*t*)) and autoregulatory (*P*_*late*_(*t*)) modules. *c* is a scaling factor between the transcriptional output of the endogenous locus and the gene regulatory function, which was measured for a transgene (***Figure 4***D). The first equation in the system shown in ***Eq. 4*** then describes the dynamics of mRNA produced from the autoregulatory element as a result of its transcriptional activity (first term on the right-hand side) and mRNA degradation (second term on the right-hand side). Further, the second equation describes the protein resulting from the autoregulatory element through the translation of the mRNA produced by this element (first term on the right-hand side) and degradation (second term on the right-hand side). Finally, as indicated in ***Eq. 4***, we assume that the autoregulatory element is only active for *t* ≥ *t*_*on*_. We note, however, that our model produced nearly identical results whether we assumed that this transition to full responsiveness occurred instantaneously at time *t*_*on*_, or whether we assumed a gradual increase in responsiveness over time (***Section S3.1***).

After time *t*_*on*_, in addition to mathematically describing the expression dynamics of the autoregulatory element, we can also model the contribution of the early element to *ftz* expression as a dynamical system. In particular, our empirical measurements of the early element transcription rate using the MCP-MS2 system revealed that its transcription rate *r*(*t*) follows an approximately exponential decay with a with a decay constant *β*^−1^ of about 21 min (***Figure 3***C; ***Figure S4***; ***Section S2.3***). We can then represent the mRNA *R*_*early*_(*t*) and protein *P*_*early*_(*t*) produced from the early module using the linear system of equations

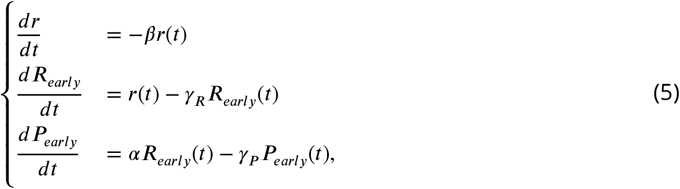

where the equation for 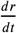 has been introduced to model the approximately exponential decay of the rate of transcription of the early element after the onset time *t*_*on*_. Here, *R*_*early*_ dynamics are dictated by the transcription rate *r*(*t*) and the mRNA degradation rate. The early protein *P*_*early*_ is determined by the translation of the mRNA stemming from the early element as well as by protein degradation. Note that, regardless of the choice of parameter values or initial conditions, *P*_*early*_(*t*) converges to 0 as *t* goes to infinity (***Section S1.2***), indicating that signaling from upstream factors is transient.

The behavior of the *ftz* regulatory system depends crucially on the quantitative values of the kinetic rates describing the molecular interactions within our model. For those parameters that were not already present in the literature, we carried out a set of experiments designed to directly measure these free parameters as described in detail in ***Section S2***. First, the gene regulatory function *f* (*P*) was measured by averaging the traces obtained from simultaneous imaging of endogenous Ftz protein and the corresponding autoregulatory response as previously shown in ***Figure 4***D. Second, the translation rate *α* was found by simultaneously measuring the rate of transcription and the resulting protein concentration in a transgenic construct where *ftz* mRNA was labeled with MS2 and the resulting Ftz protein tagged with a LlamaTag (***Figure S3***). Third, the decay in the transcription rate of the early element over time *β* was determined by fitting an exponential function to the transcriptional dynamics of the early element construct (***Figure S4***). Fourth, the scaling factor *c* was inferred by systematically comparing simulated to measured traces in the regime where the gene regulatory function was saturated and mRNA production rate was at its maximum (***Figure S5***). Finally, the mRNA and protein decay rates *γ*_*R*_ and *γ*_*P*_ were drawn from existing measurements in the literature. All parameter values are reported in ***Table S1***.

If our model of the *ftz* regulatory system is accurate, then for each nucleus along the anterior boundary of stripe 4, we should be able to predict Ftz expression state at gastrulation based only on measurements at time *t*_*on*_ = −20 min. Specifically, at this point, each nucleus will have a different initial expression rate, mRNA level, and protein level from the early module given by *r*(*t*_*on*_) = *r*_0_, *R*_*early*_(*t*_*on*_) = *R*_0_, and *P*_*early*_(*t*_*on*_) = *P*_0_, respectively. The autoregulatory element, however, will not have produced any mRNA or protein prior to -20 min, so the initial conditions for the autoregulatory module will be *R*_*late*_(*t*_*on*_) = 0 and *P*_*late*_(*t*_*on*_) = 0.

Each set of initial conditions (*r*_0_, *R*_0_, *P*_0_) for the early module defines a trajectory for *P*_*early*_(*t*) (***Figure 5***A) that will in turn drive expression from the autoregulatory module. The result is a unique overall trajectory for total Ftz concentration *P*_*total*_(*t*) = *P*_*early*_(*t*) + *P*_*late*_(*t*) (***Figure 5***B) for each nucleus. Hence, for a given set of initial conditions, we can simulate the full dynamical system starting at *t*_*on*_ and see whether total Ftz concentration exceeds a threshold at gastrulation (i.e., *P*_*early*_(*t* = 0) + *P*_*late*_(*t* = 0) > *P*_*thresh*_, where *P*_*thresh*_ is empirically determined as shown in ***Figure 2***D) in order to predict the Ftz expression state of the nucleus at gastrulation. Moreover, since increasing any one of the initial conditions can only increase total Ftz, if we plot all possible sets of initial conditions (*r*_0_, *R*_0_, *P*_0_), then a single smooth surface separates the sets that result in high Ftz at gastrulation (above the surface) from the sets that result in low Ftz at gastrulation (below the surface), as visualized in ***Figure 5***C.

**Figure 5.**
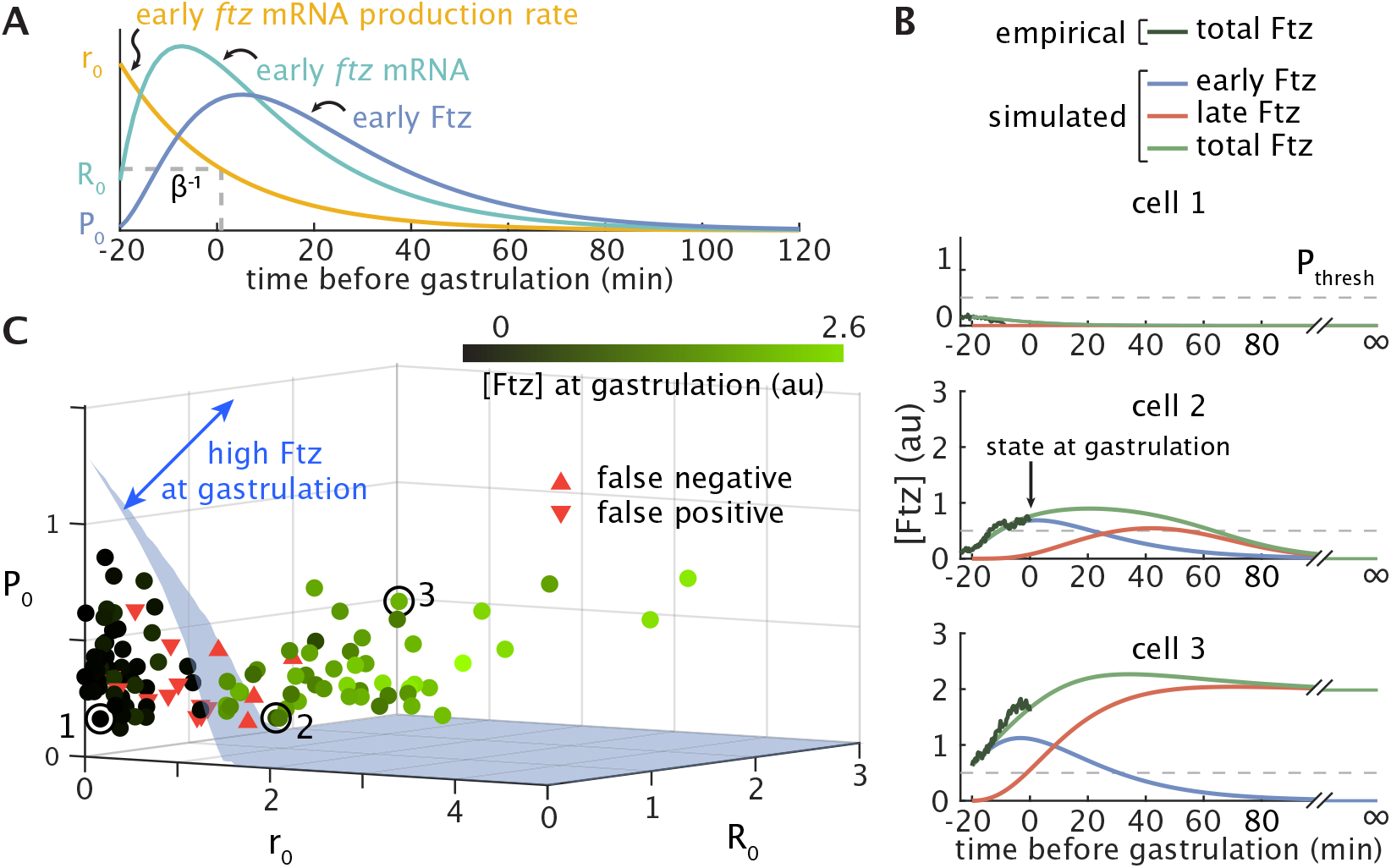
Mathematical modeling accurately predicts binary Ftz expression states at gastrulation. (**A**) Example trajectories for the mRNA production rate *r*(*t*), mRNA concentration *R*_*early*_(*t*), and protein concentration *P*_*early*_(*t*) from the early element, as described by ***Eq. 5***. (**B**) Representative traces for three nuclei comparing empirical total Ftz trajectories (dark green) to simulated trajectories (green) comprising contributions from both the early (blue) and autoregulatory (red) elements. Gray dashed line marks the experimentally determined threshold used to classify cells as high or low Ftz expression states at gastrulation (***Figure 2***D). The plotted nuclei correspond to the circled and labeled points in panel C. Note that the simulations extend well past the time that we can obtain experimental measurements. (**C**) Each nucleus has a set of initial conditions (*r*(*t*_*on*_) = *r*_0_, *R*_*early*_(*t*_*on*_) = *R*_0_, *P*_*early*_(*t*_*on*_) = *P*_0_) for the dynamics of the early element. The blue surface separates those nuclei (circles) that are predicted to express low levels of Ftz at gastrulation (below the surface) from those that are predicted to express high levels of Ftz (above the surface). Color intensity corresponds to the Ftz concentration at gastrulation. Downward-facing red triangles indicate false positives (predicted high at gastrulation, but experimentally determined to be low) and upward-facing red triangles indicate false negatives (predicted low, but experimentally determined to be high). Results are shown for *N* = 118 nuclei from 3 embryos at the anterior boundary of stripe 4, with model parameters given in ***Table S1***.

We used the smooth surface separating high and low Ftz states in our model to directly predict Ftz state at gastrulation for our experimentally measured nuclei. To avoid conflating measurements with predictions, we analyzed a set of *N* = 3 embryos independently from those used to obtain all model parameter values, including the gene regulatory function (***Section S2***). Our model achieved 86.4% binary classification accuracy (102 of 118 nuclei) based on the system’s initial condition at *t* = *t*_*on*_ relative to the experimentally measured Ftz expression state at gastrulation (*t* = 0) determined by thresholding measured Ftz concentrations for each nucleus. The majority of classification errors were false negatives derived from a single embryo; excluding this embryo from analysis produced an overall prediction accuracy of 93.5% (72 of 77 nuclei). For the two well-predicted embryos, the empirical false negative, false positive, and total error rates aligned extremely well with error rates derived from stochastic simulations (***Section S3.4***). As a result, it is plausible that many of the remaining classification errors can be attributed to noisy gene expression dynamics.

Taken together, these observations suggest that our simple model captures the essential deterministic components of Ftz dynamics and is able to predict Ftz expression state at gastrulation from knowledge of the initial conditions of the early element. However, being able to predict Ftz expression state at gastrulation does not guarantee that the autoregulatory element permanently locks the Ftz expression fate for the rest of development through bistability. To determine whether such developmental memory is at play, we need to further analyze our theoretical model.

### 2.4 The *ftz* autoregulatory module is bistable and remembers binary cell state

Having established that our model accurately predicts transient Ftz expression state at gastrulation, we next analyzed the behavior of the early and autoregulatory modules separately, with the goal of uncovering whether the autoregulatory module is bistable and, as a result, retains a long-term binary memory of these Ftz levels.

We first performed a test of the autoregulatory module to ascertain whether it is bistable in the absence of Ftz contribution from the early element (*P*_*early*_(*t*) = 0). Such bistability would enable the module to maintain a high or low expression state even once upstream regulatory factors binding the early element have degraded. Following the procedure described in ***Box 1*** and ***Section S1.3***, we identified the steady states where mRNA and protein concentrations no longer change in time. Our analysis revealed that, indeed, the empirically determined model parameters set the autoregulatory module in a bistable regime (***Figure 6***A) characterized by the presence of two stable steady states corresponding to low and high Ftz values. This result indicates that the *ftz* autoregulatory module is capable of maintaining high or low levels of *ftz* expression indefinitely.

**Figure 6.**
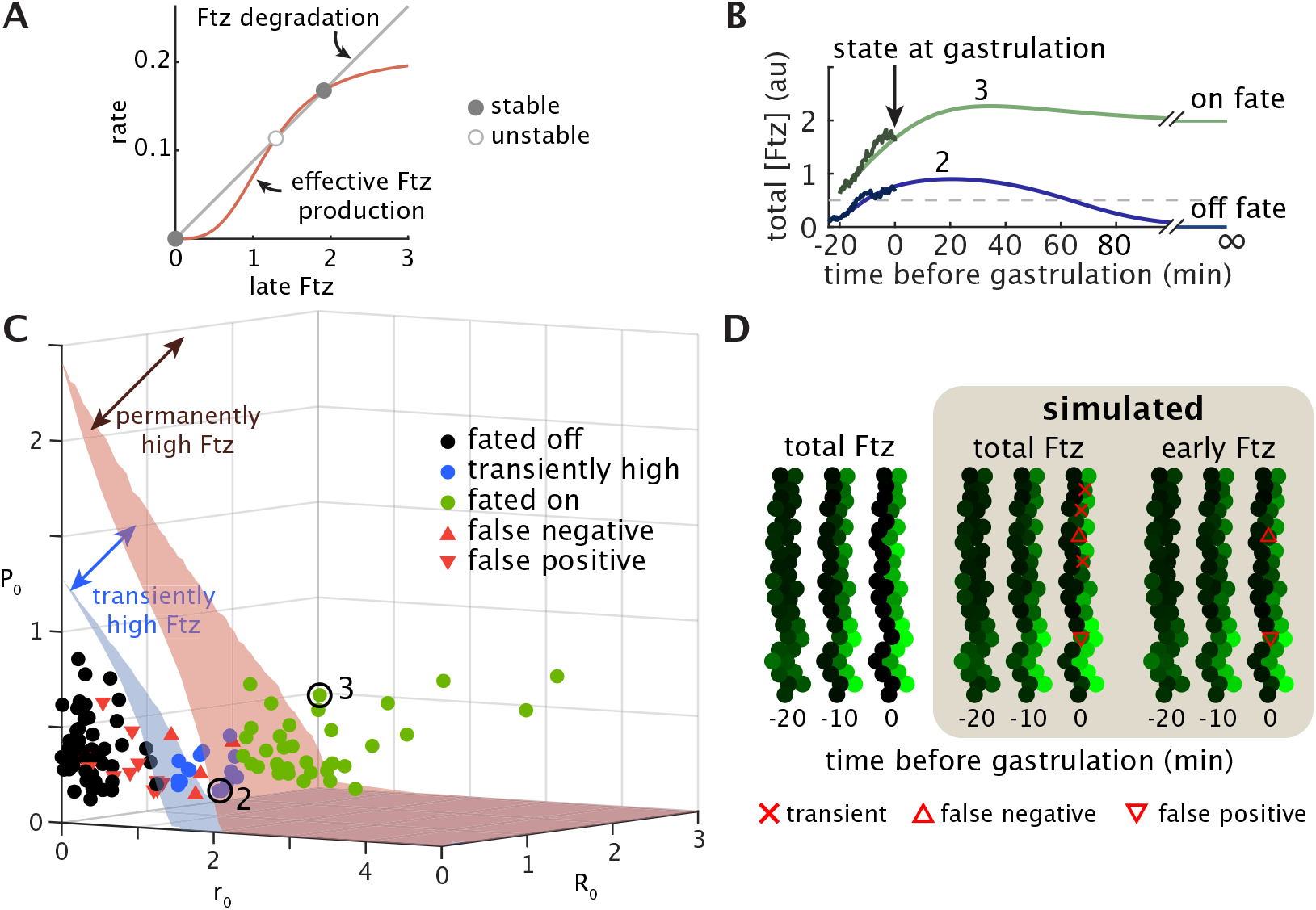
Quantitative mathematical model reveals that the *ftz* autoregulatory module is bistable. (**A**) The intersection between Ftz degradation rate and the effective Ftz production rate (i.e., late Ftz production rate adjusted for late *ftz* mRNA production and decay rates) reveals that the autoregulatory module is bistable, as described in ***Box 1*** and ***Section S1.3***. (**B**) Most cells that express high Ftz state at gastrulation reach the high fate at steady state (cell 3), but some transiently express high Ftz at gastrulation before ultimately reaching the low fate (cell 2). The threshold (dashed gray line) is only used to determine state at gastrulation; fate is decided by which of the two stable steady states is approached by the system in simulation at long times. (**C**) The early module dictates the transient dynamics of Ftz as well as the fate it is predicted to adopt at infinite time once the autoregulatory element has reached steady state. The blue surface is repeated from ***Figure 5*** and separates nuclei into low (below surface) and high (above surface) Ftz state at gastrulation. The red surface (switching separatrix) separates nuclei predicted to adopt the low Ftz fate (black circles) from those predicted to adopt the high Ftz fate (green circles), where by “fate” we mean expression level at infinite time (steady state). Nuclei between the blue and red surface (blue circles) are considered transiently high in that they express high Ftz at gastrulation but are predicted to adopt the low Ftz fate. Data are plotted for *N* = 118 nuclei from 3 embryos. Red triangles indicate false negatives (upward) and false positives (downward) as determined from classification at gastrulation. (**D**) Results from a representative embryo show that the experimentally measured stripe pattern (left) is recapitulated by simulation (middle). A stripe pattern is still evident at gastrulation even from the predicted early Ftz concentration alone (right). Nuclear intensities at all time points are normalized to the predicted steady-state high Ftz concentration. Red “x”s denote nuclei with a transiently high Ftz state at gastrulation; triangles denote false positives (downward) and false negatives (upward). Parameters for simulation are as given in ***Table S1***.

Since our model predicts that the autoregulatory element is capable of remembering Ftz expression levels, the question thus arises of whether the module actually becomes responsive to Ftz concentration in time to lock the transient state observed in the blastoderm into a permanent cell fate. Otherwise, if the bistable module becomes responsive too late, cells could transiently express high Ftz at gastrulation without committing to stably expressing high Ftz in the long term.

To distinguish between these two scenarios, we compared the binary classification of nuclei into high or low Ftz expression states at gastrulation as introduced in the previous section with the final high or low expression fate predicted by the model in steady state (***Figure 6***B). Similarly to the case of binary classification, the regions of parameter space (*r*_0_, *R*_0_, *P*_0_) resulting in high fates are separated from the region of parameters resulting in low fates by a surface called the switching separatrix (***Sootla et al., 2016***), so named because if the initial conditions of a cell are above the surface, then the bistable autoregulatory module will switch on. The initial conditions for all nuclei at the anterior of stripe 4 are plotted in ***Figure 6***C alongside the switching separatrix (red surface) and the surface for transient binary classification of Ftz state at gastrulation (blue surface).

If we restrict our analysis to the “best predicted” nuclei (expression state correctly classified at gastrulation and with relatively low cumulative error between simulation and empirical measurement over time; see ***Section S3.1***), 93.7% (74 of 79 nuclei) were predicted to maintain their binary state and adopt the corresponding Ftz expression fate in steady state, as exemplified by cell 3 in ***Figure 6***B. The remaining 6.3% of nuclei (5 of 79), despite having a high Ftz expression state at gastrulation, were predicted to drop to a low Ftz fate in steady state as shown by cell 2 in ***Figure 6***B. No nuclei classified as low Ftz expression state at gastrulation were predicted to express high Ftz after gastrulation. Thus, the autoregulatory element ensures that the vast majority of cells adopt a fate matching the transient state at gastrulation.

While it is clear that the autoregulatory element establishes developmental memory by fixing the Ftz expression fate in steady state, we wondered whether this memory was already at play at gastrulation, or whether the early module is principally responsible for setting Ftz state at gastrulation. Our simulations show that a stripe pattern is already evident at gastrulation from the contribution of the early protein alone, ignoring the autoregulatory contribution (***Figure 6***D). Thus, it appears that the anterior boundary of stripe 4 at gastrulation is defined by the regulatory activity of upstream factors binding the early element in a manner that is largely independent of the activity of the autoregulatory element. Therefore, this result supports the conclusion that the autoregulatory module is bistable and that it primarily serves to commit cells to fates predetermined by the early element.

### 2.5 Ftz fate is robustly specified in half an hour

Given that the *ftz* autoregulatory element acts as a memory module, we might ask how long it takes nuclei to convert the transient expression state of the early module into a stable cellular memory, thereby establishing an expression fate at long times beyond gastrulation. We posit that this timespan corresponds to the classical notion of a commitment window, defined as the period of time during which a cell integrates information from external factors to decide its fate (***Dalton, 2015***; ***McNeely and Dwyer, 2021***). It can be difficult to access temporal features of development such as the commitment window *in vivo*, in part due to the technical challenge of measuring and systematically manipulating input signals while simultaneously monitoring the resulting gene expression programs in individual cells within intact tissues or organisms (***Bending et al., 2018***; ***Johnson and Toettcher, 2018***).

Our mathematical model provided us with a unique opportunity to examine the commitment window by altering the timing of developmental events *in silico*. Because the low Ftz fate at steady state is the default expression state—the autoregulatory module will always produce zero protein in the absence of a transient signal from the early module—the commitment window primarily determines whether the cell has enough time to detect if transient Ftz concentrations are high and, if so, to adopt a trajectory destined for a high steady-state expression fate. Thus, for the results reported in this section, we restricted our analysis to a subset of the best predicted nuclei, the nuclei that were correctly classified at gastrulation and have relatively low cumulative error between simulation and experiment over time (***Section S3.1***), that were also predicted to adopt the high fate according to the switching separatrix analysis (*N* = 21; ***Figure 6***B).

We define the commitment window as *t*_*off*_ − *t*_*on*_, where *t*_*on*_ indicates the start of autoregulatory responsiveness and *t*_*off*_ is the time when upstream factors stop controlling the transcriptional dynamics of the early element (***Figure 7***A and B). In other words, the commitment window represents the total amount of time during which both the early and autoregulatory modules dictate *ftz* expression (***Figure 7***C), and serves as an estimate for how long the autoregulatory module has to establish a memory of transient Ftz state. We define *t*_*off*_ such that *r*(*t* ≥ *t*_*off*_) = 0, where *r*(*t*) is the mRNA production rate of the early element. Before *t*_*off*_, *r*(*t*) follows the usual exponential decay with rate 1/*β* = 21 min (***Figure 7***B).

**Figure 7.**
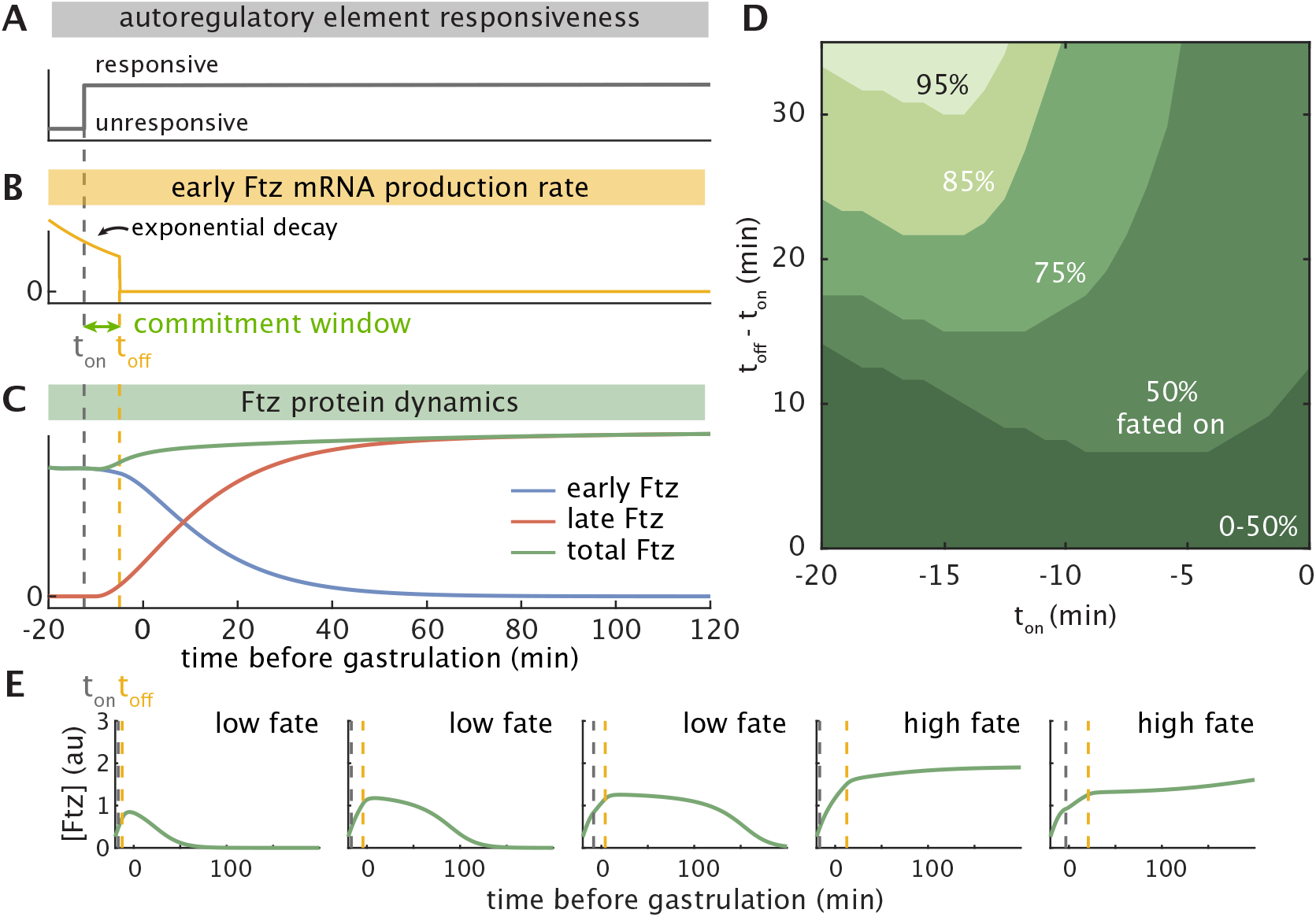
*In silico* analysis reveals the window of commitment of cells to the high Ftz fate. (**A-B**) Schematic illustrating (A) *t*_*on*_, the time the autoregulatory element becomes responsive, and (B) *t*_*off*_, the time the early element ceases production. The commitment window *t*_*off*_ − *t*_*on*_ is the timespan during which both upstream factors and autoactivation dictate Ftz expression, and serves as an estimate for how long the autoregulatory module has to establish a memory of the signal from the early module. (**C**) Simulated Ftz concentrations resulting from the timing pictured in A and B. (**D**) Contour plot showing what percentage of the subset of analyzed nuclei (see text) reach high Ftz fate under wild-type conditions (*N* = 21; ***Figure 6***B, ***Section S3.1***) still reach that fate as the commitment window *t*_*off*_ − *t*_*on*_ is varied and *t*_*on*_ is delayed relative to the measured -20 min (see text). (**E**) Simulated single-nucleus traces of total Ftz for varying values of *t*_*on*_ and the commitment window can cause the transient dynamics to vary quite dramatically. Dashed lines denote the *t*_*off*_ (gold) and *t*_*on*_ (gray) values used for each plot. Parameters are as in ***Table S1***. See also ***Figure S10***.

To determine how long of a commitment window allows cells to convert transiently high Ftz concentrations into permanently high Ftz fates, we asked whether our analyzed subset of nuclei still reached the high fate as we systematically varied the commitment window. We solved the dynamical system from ***Eq. 4*** and ***Eq. 5*** with commitment windows of increasing length and start times *t*_*on*_ between -20 min and 0 min and recorded which nuclei reached the high Ftz fate. Our results are reported in terms of the fraction of nuclei within the subset that adopted the high Ftz fate for each timing condition.

***Figure 7***D shows the fraction of nuclei predicted to adopt the high Ftz fate as a function of the commitment window and *t*_*on*_. For example, assuming *t*_*on*_ ≤ −13 min, a commitment window of 34 min results in 95% of nuclei achieving high steady-state Ftz levels without further signaling from the early element (***Figure 7***D). Our results also reveal that there is a gradual dropoff in the fraction of cells that do not commit to the high fate as the commitment window is shortened or *t*_*on*_ is delayed. This suggests that slight temporal perturbations during development are unlikely to cause catastrophic patterning failures in which all cells suddenly adopt the low Ftz fate. Rather, we might expect small changes in timing to affect the fates of only a small percentage of cells.

Our analysis of the commitment window indicates that about half an hour can suffice for the vast majority of high-fated cells to stably commit to that fate. Proper fate specification, however, does not guarantee similarity in the temporal trajectories of Ftz concentration, as evidenced by the wide range of dynamics observed in simulated traces for varying *t*_*on*_ and *t*_*off*_ with the same initial conditions for the early module (***Figure 7***E; ***Figure S10***). Thus, if the transient Ftz trajectory, not just its ultimate fate, is instructive for downstream genes, then the need for proper regulation of these genes may place stricter constraints on the relative timing of the early and autoregulatory elements than those that are imposed by the specification of steady state Ftz fate alone.

## 3 Discussion

For decades, developmental biologists have used the concept of Waddington’s landscape to conceptualize the adoption of discrete cell fates. Under this framework, cells roll down valleys in a predetermined landscape to adopt their ultimate fates. This framework has been repeatedly mathematicized using dynamical systems theory (***Wang et al., 2011***; ***Furusawa and Kaneko, 2012***; ***Jaeger and Monk, 2014***; ***Corson and Siggia, 2017***; ***Sáez et al., 2022***). Many of these studies have hypothesized that positive autoregulation (***Crews and Pearson, 2009***) helps establish and maintain binary cell fates through bistability (***Zernicka-Goetz et al., 2009***; ***Soldatov et al., 2019***), which can be thought of as introducing forks in Waddington’s landscape. Though experiments in cell culture and fixed tissue have provided evidence for the bistability of various autoregulatory modules found within gene regulatory networks, until now, these results have not been confirmed by direct examination of dynamics in intact, living embryos.

In this work, we utilized live imaging approaches to quantitatively characterize the dynamics of the fruit fly *ftz* regulatory system *in vivo*. We elucidated tight temporal coordination between the two enhancer elements that regulate *ftz* expression (***Figure 3***), and combined dynamical systems modeling with biophysical measurements to show that the bistability of the autoregulatory module can remember otherwise transient expression levels driven by upstream factors (***Figure 6***). Based on the prevalence of autoregulatory motifs found in nature (***Alon, 2006***; ***Peter and Davidson, 2015***), we speculate that the approach employed by the Ftz system to decide cell fate is not limited to fruit flies, but might also be widely adopted during development in other organisms.

One of our central discoveries is that *ftz* autoregulation is triggered at a specific developmental time rather than by a threshold concentration of Ftz protein driven by the early element. Recent work has suggested candidates for “timer genes” that are expressed at distinct developmental time points and appear to facilitate the expression of other genes (***Clark and Akam, 2016***; ***Clark and Peel, 2018***; ***Clark et al., 2022***). At least one of these timer genes, Odd-paired, has been shown to exhibit chromatin opening activity (***Soluri et al., 2020***), and two, Caudal and Dichaete, have been shown to bind directly to the *eve* autoregulatory element (***MacArthur et al., 2009***). We speculate that timer genes might also bind the *ftz* autoregulatory element to trigger its responsiveness to Ftz protein. Furthermore, the tight temporal coordination we observed between the transcriptional activity driven by the early and autoregulatory elements suggests that sequentially expressed timer genes might differentially regulate enhancers to precisely coordinate gene expression during development.

A basic assumption of our work is that genetic networks exhibit modularity (***Hartwell et al., 1999***; ***Bolouri and Davidson, 2002***; ***Del Vecchio et al., 2016***), meaning the network can be broken down into parts (modules), each with some inputs and outputs connected to other modules. The behavior of the whole network can then be predicted from the behavior of the modules in isolation. How to define the modules, or equivalently how to break the network down into parts, has been the subject of much discussion (***Hartwell et al., 1999***). As we have argued in this work, topology, the pattern of interactions among molecular species, is not enough to determine the behavior of a module. Indeed, its dynamics will depend on the values of the parameter governing these interactions (***Angeli et al., 2004***; ***Ingram et al., 2006***; ***Graham et al., 2010***; ***Blanchini and Franco, 2014***; ***Khammash, 2016***; ***Verd et al., 2019***).

There is inherent flexibility in deciding which physical components to include in a module, and hence which quantities act as parameters and which as inputs to the module. For example, for the *ftz* autoregulatory system, the distinction between the early and autoregulatory elements led us to structure the autoregulatory module with input *P*_*early*_ and output *P*_*late*_ ***Figure S2***. This is not the only way to define the module; notably, if we knew which regulatory factors rendered the autoregulatory element responsive, then we could include those as inputs and modify the model accordingly.

Our fairly simple representation of the autoregulatory module can predict the fate of *ftz* expression from arbitrary trajectories of early Ftz. As a result, we can predict the effect of modifications to upstream signaling on the resulting gene expression patterns. This, in turn, allows us to ask what forms of input are appropriate to achieve particular patterning outcomes. Such ability to reverse engineer the process of cellular decision-making could facilitate designing perturbations to manipulate the system, identifying constraints placed on upstream modules by the needs of downstream modules, and analyzing whether biologically evolved signals match those that are mathematically “optimal” for such needs as speed of patterning (***Pezzotta and Briscoe, 2022***) or information transmission (***Tkačik and Gregor, 2021***). Different methods of generating predictions may be appropriate depending on the types of inputs under consideration. In this paper, the fact that increasing any one of the parameters that define the early Ftz input (*r*_0_, *R*_0_, *P*_0_) increases total Ftz concentration at all points in time (a property known as monotonicity; ***Angeli et al. (2004***); ***Sootla et al. (2016***)) made it possible to analyze our model using a switching separatrix. However, this may not be true for other regulatory systems, as in the case where a gene within a module represses its own production.

Throughout developmental biology, the concept of a commitment window has been repeatedly utilized to describe the amount of time cells need to be exposed to upstream signals in order to decide their developmental fates (***Dalton, 2015***; ***McNeely and Dwyer, 2021***). Our quantitative dynamical systems model made it possible to conduct a detailed examination of this commitment window, and to identify what fraction of cells adopt certain fates as developmental timing is varied. From an engineering perspective, we may consider a gene expression pattern as a “design specification” that must be achieved with a prescribed level of precision, and work backwards to see what inputs satisfy this requirement. Our approach complements existing work on precision that emphasizes how tightly protein concentrations are controlled (***Gregor et al., 2007***) and how accurately cells can locate their position by reading out concentrations of upstream factors (***Dubuis et al., 2013***; ***Petkova et al., 2019***). In particular, the latter approaches indicate what level of precision is actually achieved by a patterning network, while our framing focuses rather on what range of parameters allow a system to attain a predefined level of precision. A combination of the two perspectives could help elucidate what biophysical and evolutionary factors influence stochastic variation in phenotypes, including how precise expression patterns must actually be to produce functional, healthy organisms. In any case, we hope our results act as one more step toward grounding robustness or reproducibility in developmental patterning on a quantitative, probabilistic level.

In summary, by turning widespread schematic models of autoactivation modules into precise mathematical statements and experimentally testing the resulting predictions, we have provided support for a widely held hypothesis about how developmental fates are established in embryos. In the future, combining quantitative measurements with precise spatiotemporal perturbations (***Goglia and Toettcher, 2019***) and synthetic reconstitution methods (***McNamara et al., 2022***) promises to enable yet another iteration of the dialogue between theory and experiment that constitutes the basis of our work, ultimately leading to a predictive understanding of function in developmental networks and the myriad forms and fates to which they give rise.

## Methods and Materials

### Cloning and Transgenesis

The fly lines used in this study were generated by inserting transgenic reporters into the fly genome or by CRISPR-Cas9 genome editing, as described below. See ***Table S2*** for detailed information on the plasmid sequences used in this study.

#### Creation of tagged *fushi tarazu* (*ftz*) gene using CRISPR-Cas9

Ftz-EGFP-LlamaTag fusion design is based on previously published transgenic line (***Bothma et al., 2018***). To tag endogenous *ftz* locus with EGFP-LlamaTag, we used CRISPR-mediated homology-directed repair with donor plasmid synthesized by Genscript. gRNA was designed using the target finder tool from flyCRISPR (https://flycrispr.org), and cloned based on the protocol from ***Gratz et al. (2015***). yw;nos-Cas9(II-attP40) transgenic line was used as the genomic source for Cas9 and the embryos were injected and screened by BestGene Inc.

#### Creation of *ftz* autoregulatory element reporter

The *ftz* autoregulatory element sequence is based on 4.4kb DNA segment described in (***Hiromi et al., 1985***). The *ftz* autoregulatory element reporter was constructed by combining the enhancer sequence with an array of 24 MS2 stem loops fused to the *D. melanogaster yellow* gene (***Bothma et al., 2018***). The construct was synthesized by Genscript and injected by BestGene Inc into *D. melanogaster* embryos with a ΦC31 insertion site in chromosome 3 (Bloomington stock #9750; landing site VK00033; cytological location 65B2).

#### Transgenes expressing EGFP and MCP-mCherry

The fly line maternally expressing MCP-mCherry (chromosome 3) was constructed as described in (***Bothma et al., 2018***). The fly line maternally expressing vasa-EGFP (chromosome 2) was constructed as described in (***Kim et al., 2021***). To simultaneously image protein dynamics using LlamaTags and transcription using MCP-MS2 system, we combined the vasa-EGFP transgene with MCP-mCherry to construct a new line (yw;vasa-EGFP;MCP-mCherry) that maternally expresses both proteins.

### Fly lines

To measure Ftz transcription and protein levels simultaneously, we performed crosses to generate virgins carrying transgenes that drive maternal EGFP, MCP-mCherry, the LlamaTagged Ftz locus along with *ftz* autoregulatory element reporter (yw; 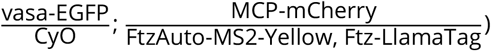. These flies were then crossed with males having both the *ftz* autoregulatory element reporter and the LlamaTagged Ftz locus (yw; +; FtzAuto-MS2-Yellow, Ftz-LlamaTag). This resulted in the embryo carrying maternally deposited EGFP, MCP-mCherry, and two copies of the LlamaTagged Ftz locus and *ftz* autoregulatory element reporter.

### Embryo preparation and data collection

The embryos were prepared following procedures described in (***Garcia et al., 2013***; ***Bothma et al., 2018***; ***Lammers et al., 2020***). Embryos were collected and mounted in halocarbon oil 27 between a semipermeable membrane (Lumox film, Starstedt, Germany) and a coverslip. Confocal imaging on a Zeiss LSM 780 microscope was performed using a Plan-Apochromat 40x /1.4NA oil immersion objective. EGFP and MCP-mCherry were excited with laser wavelengths of 488 nm (25.0 *μ*W laser power) and 594 nm (15.0 *μ*W laser power), respectively. Fluorescence was detected using the Zeiss QUASAR detection unit. Image resolution was 512 × 512 pixels, with pixel size of 0.231 *μ*m. Sequential z-stacks separated by 0.5 *μ*m were acquired. Specimens were imaged from mid nuclear cycle 14 until the start of gastrulation.

### Image processing

Image analysis of live embryo movies was performed based on the protocol in (***Garcia et al., 2013***; ***Reimer et al., 2021***), which included nuclear segmentation, spot segmentation, and tracking. In addition, the nuclear fluorescence of Ftz was calculated based on a nuclear mask generated from the MCP-mCherry channel. Ftz concentration for individual nuclei was extracted based on the integrated amount from maximum projection along the z-stack. The GFP background was calculated based on a control experiment and subsequently subtracted from the data.

### Numerical analysis and simulations

Numerical analysis and simulations were carried out using custom scripts in MATLAB (2017b). The switching separatrix and analogous surface for transient Ftz state at gastrulation were estimated using a modification of the algorithm in ***Sootla et al. (2016***), which employs a combination of bisection and random sampling to estimate the upper and lower bounds of the separatrix surface. Stochastic differential equations were simulated using the Euler-Maruyama method. More detailed descriptions of the procedures are available in the Supplementary Text.

## Acknowledgments

MLP was supported by the European Molecular Biology Laboratory Interdisciplinary Postdoc Programme (EIPOD4 fellowships), cofunded by Marie Skłodowska-Curie Actions (grant agreement number 847543). HGG was supported by the Burroughs Wellcome Fund Career Award at the Scientific Interface, the Sloan Research Foundation, the Human Frontiers Science Program, the Searle Scholars Program, the Shurl and Kay Curci Foundation, the Hellman Foundation, the NIH Director’s New Innovator Award (DP2 OD024541-01), NSF CAREER Award (1652236), an NIH R01 Award (R01GM139913) and the Koret-UC Berkeley-Tel Aviv University Initiative in Computational Biology and Bioinformatics. HGG is also a Chan Zuckerberg Biohub Investigator.

## Author contributions

Conceptualization: JZ, MLP, HGG

Methodology: JZ, MLP, HGG

Resources: JZ, MLP, MN, HGG

Investigation: JZ, MLP, HGG

Visualization: JZ, MLP, HGG

Funding acquisition: HGG

Project administration: HGG

Supervision: HGG

Writing – original draft: JZ, MLP, HGG

Writing – review & editing: JZ, MLP, HGG

## Data and materials availability

All materials are available upon request. All data in the main text or supplementary materials are available upon request. All code is available in this paper’s github repository.

## Supplementary Information

### Supplementary Figures

**Figure S1.**
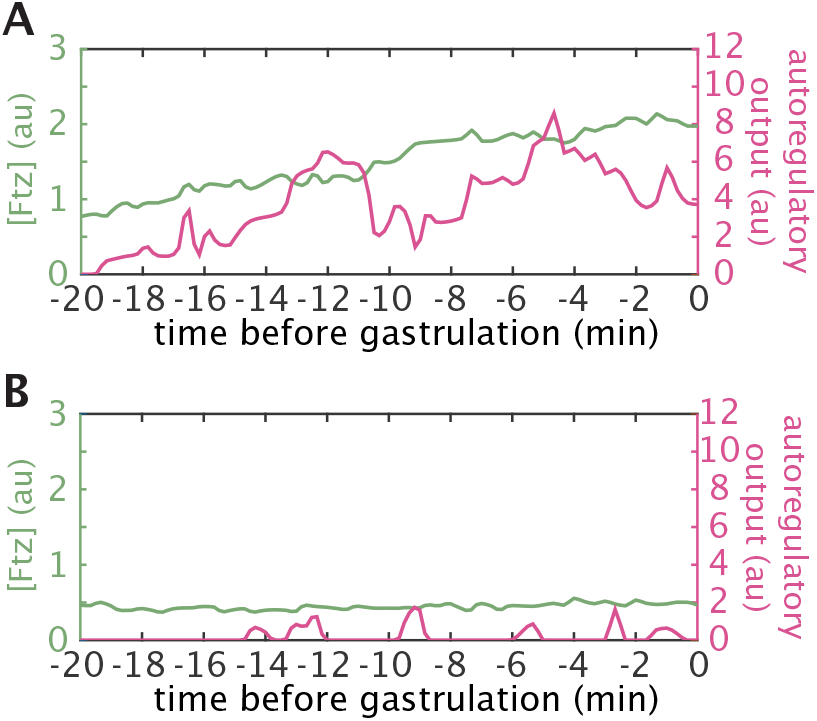
Single-cell traces of Ftz protein and autoregulatory activity show that high or low Ftz concentration results in distinct temporal dynamics. Green and magenta lines correspond to Ftz protein and transcriptional activity of the autoregulatory element, respectively, obtained as described in fig:autoregulationA. (**A**) High level of Ftz protein results in continued active transcription from the autoregulatory element. (**B**) Low level of Ftz protein results in a much weaker autoregulatory response.

**Figure S2.**
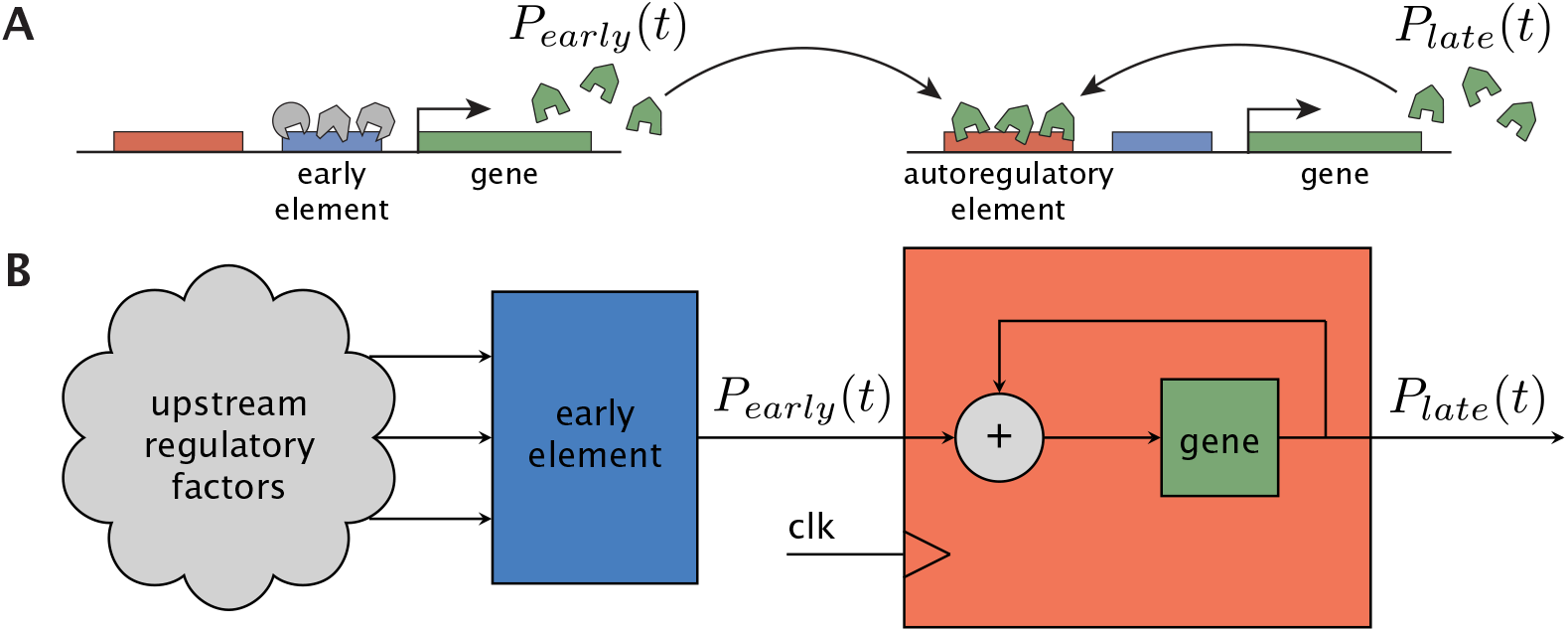
Two ways to visualize a genetic network as an interconnection of modules. (**A**) Schematic of Ftz regulation in which populations of Ftz protein are differentially labeled based on whether they are generated by regulation from the early element (*P*_*early*_(*t*)) or from the autoregulatory element (*P*_*late*_(*t*)). The two populations are summed to generate the total Ftz concentration that binds the autoregulatory element. (**B**) The Ftz regulatory system can be visualized as a block diagram to highlight how a modular network decomposition might be inspired by similar breakdowns of electrical circuits and other engineering control systems (***Cosentino and Bates, 2011***). Here, unspecified upstream factors regulate the early element module, which produces output *P*_*early*_(*t*). This, in turn, acts as the input to the autoregulatory module, which has been schematized as a latch with a clock signal (clk) to show that the element only becomes responsive at a particular time. The output *P*_*late*_(*t*) can be summed with *P*_*early*_(*t*) to recover total Ftz concentration. Relative to the more traditional schematic in (A), this representation includes dynamic signals (*P*_*early*_(*t*) and *P*_*late*_(*t*)), a timing element that triggers autoregulatory responsiveness, and a clear separation of inputs and outputs from each module.

### Supplementary Tables

**Table S1.** Quantitative parameters used throughout this work.

| Parameter | Description | Source | Value | Units |
| --- | --- | --- | --- | --- |
| $\alpha$ | translation rate | this work | $0.082 \pm 0.004$ | protein AU (mRNA AU min)<sup>-1</sup> |
| $\tau_R$ | mRNA half-life | (Edgar et al., 1986) | 7 | min |
| $\gamma_R$ | mRNA decay rate | $\frac{\ln 2}{\tau_R}$ | 0.0990 | min<sup>-1</sup> |
| $\tau_P$ | protein half-life | (Bothma et al., 2018) | 7.9 | min |
| $\gamma_P$ | protein decay rate | $\frac{\ln 2}{\tau_P}$ | 0.0877 | min<sup>-1</sup> |
| $f(P)$ | autoactivation gene regulatory function | this work | $\frac{aP^n}{K^n + P^n}$ | mRNA AU (protein AU)<sup>-1</sup> |
| $a$ | maximum transcription rate | this work | $5.555 \times 10^5$ | mRNA AU (protein AU)<sup>-1</sup> |
| $K$ | - | this work | $1.216 \times 10^6$ | protein AU |
| $n$ | Hill coefficient | this work | 3.264 | (none) |
| $c$ | ratio of transcription rate endogenous to transgene | this work | $0.45 \pm 0.02$ | (none) |
| $\beta$ | decay rate of early mRNA production | this work | $0.0479 \pm 0.0021$ | min<sup>-1</sup> |

**Table S2.** List of plasmids used in this study.

| Name | Function |
| --- | --- |
| pBPhi-FtzEarly-MS2 | *ftz* early element MS2 reporter |
| pBPhi-FtzUpstream-MS2-Yellow | *ftz* autoregulatory element MS2 reporter |
| pUC57-Ftz-LlamaTag-dsRed | Donor plasmid for Ftz-LlamaTag CRISPR knock-in fusion |
| pU6-3-gRNA-Ftz-1 | guide RNA 1 for Ftz-LlamaTag CRISPR knock-in fusion |
| pU6-3-gRNA-Ftz-2 | guide RNA 2 for Ftz-LlamaTag CRISPR knock-in fusion |
| pU6-3-gRNA-Ftz-3 | guide RNA 3 for Ftz-LlamaTag CRISPR knock-in fusion |

### Supplementary Movie

**Movie S1**. Two-color imaging of input Ftz protein concentration and the output autoregulatory dynamics at stripes 3, 4 and 5.

### Supplementary Text

#### S1 Modeling

##### S1.1 Autoregulatory element

We use *R*_*early*_(*t*) to describe the concentration of *ftz* mRNA transcribed from the early element, which is translated into protein *P*_*early*_(*t*). We define *R*_*late*_(*t*) as the ftz mRNA transcribed from the autoregulatory element and translated into protein *P*_*late*_(*t*). The total Ftz protein in the cell at time *t* is given by *P*_*total*_(*t*) = *P*_*early*_(*t*) + *P*_*late*_(*t*). The dynamical equations describing the temporal evolution of mRNA and protein are

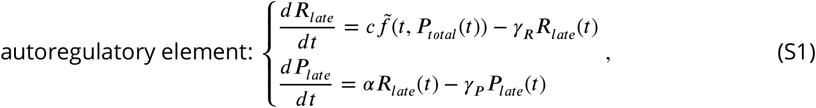

where *γ*_*R*_ and *γ*_*P*_ are the decay rates of mRNA and protein respectively, *α* is the translation rate, and *c* is a scaling factor equivalent to the ratio of maximum production rate from the endogenous locus vs. the transgene (where 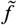 is measured). Note that 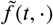, the gene regulatory function for the autoregulatory element, is time dependent. Unless otherwise stated, we will assume

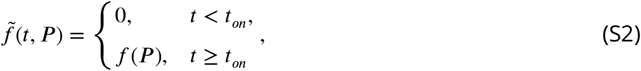

where

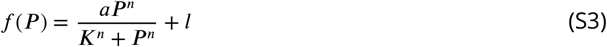

is the sigmoid describing the autoregulatory relationship at maximum amplitude. Using the definition of *P*_*total*_(*t*) we will then write ***Eq. S1*** as

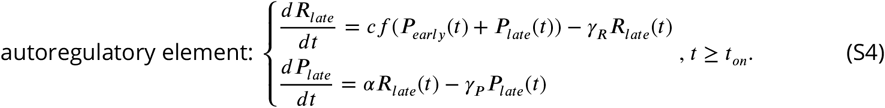

Thus, *P*_*early*_(*t*) acts as the sole time-varying input to the autoregulatory element. Since autoregulation does not begin until time *t*_*on*_, the initial conditions for ***Eq. S4*** are fixed at *R*_*late*_(*t*_*on*_) = 0 and *P*_*late*_(*t*_*on*_) = 0.

##### S1.2 Early element

From our empirical measurements, we observed that the production rate *r*(*t*) of early *ftz* mRNA is well approximated by an exponential decay (***Figure S4***), allowing us to model *R*_*early*_(*t*) and *P*_*early*_(*t*) through the dynamical system

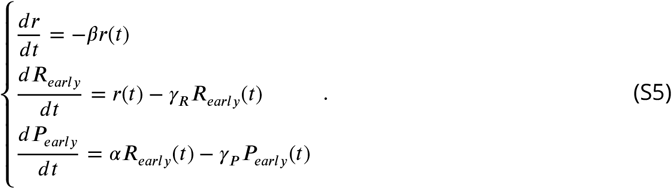

As before, *α* is the translation rate and *γ*_*R*_ and *γ*_*P*_ are the decay rates of mRNA and protein.

Because ***Eq. S5*** is linear, it can be equivalently written as

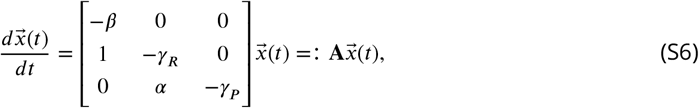

where

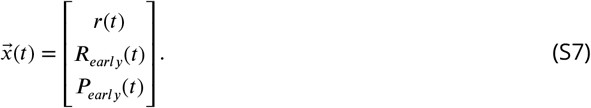

The analytical solution is then given by

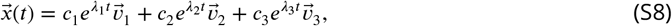

where *λ*_1_, *λ*_2_, *λ*_3_ are the eigenvalues of *A* corresponding to eigenvectors 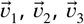 respectively. In particular,

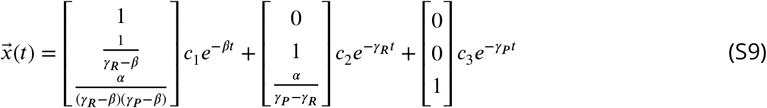

where

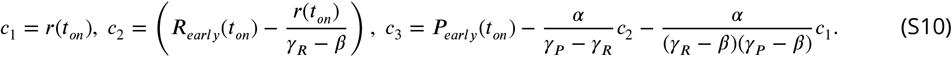

Thus, the input *P*_*early*_(*t*) to the autoregulatory element is completely characterized by three parameters (*r*(*t*_*on*_), *R*_*early*_(*t*_*on*_), *P*_*early*_(*t*_*on*_)) corresponding to the initial conditions for ***Eq. S5***. Since every term in the solution is multiplied by an exponential that decays in time, all state variables will tend to 0 as *t* goes to infinity.

##### S1.3 Intersection test for bistability

In ***Box 1*** in the main text we give an example of how to identify stable steady states in a model that only acknowledges protein concentration and ignores mRNA dynamics. Here, we show how to use the graphical intersection test on systems including both mRNA and protein.

To reiterate, our dynamical system is described by ***Eq. S4*** is given by

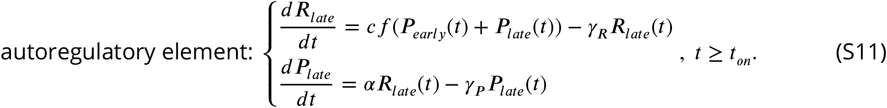

Steady states are system states at infinite time. Since we know that *P*_*early*_(*t*) goes to 0 at long times, the steady states for ***Eq. S4*** are equivalent to the steady states of

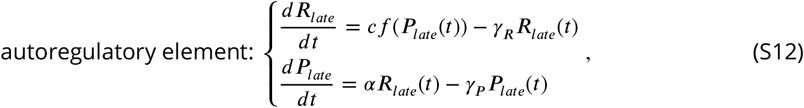

which has no input and can therefore be analyzed for steady states by standard methods.

By definition, at steady state the derivative of the different molecules species with respect to time equals zero, allowing us to write

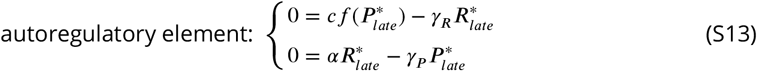

where 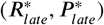 is a steady state. Then, we rearrange the bottom equation to get

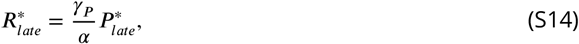

which we plug into the top equation to get

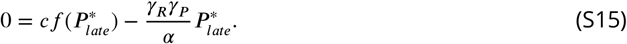

From here, we rearrange terms to recover

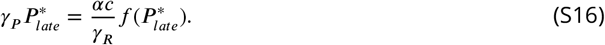

Hence, the intersections of a line of slope *γ*_*P*_ with the right-hand side give the steady-state late protein concentrations 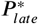, from which we can recover 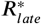 through ***Eq. S14***. For a plot of the intersection test, see ***Figure 6***A in the main text.

#### S2 Parameter estimation

##### S2.1 Regulatory function *f* (*P*) of the *ftz* autoregulatory element

We calculated the regulatory function (*f* (*P*)) of the *ftz* autoregulatory element (shown in ***Figure 4***D) for the anterior boundary of stripe 4. To make this possible, we identified the boundary in a manually selected image frame prior to gastrulation by extracting two adjacent columns of cells, each corresponding to high or low Ftz concentration. For each cell, we obtained the MS2 signal, which is a proxy for the instantaneous rate of transcription (***Garcia et al., 2013***; ***Bothma et al., 2014***; ***Lammers et al., 2020***), and the Ftz fluorescence for each time point. Next, we binned data points within a specific temporal window into ten quantiles. We averaged the MS2 and Ftz signals belonging to the same quantile, then fit a Hill function to the resulting values to obtain the regulatory function of the *ftz* autoregulatory element within that time window. We repeated the process in *N* = 7 embryos and pooled data from all embryos for subsequent analysis over four temporal windows (−20 to -15, -15 to -10, -10 to -5, and -5 to 0 min) to obtain the trend shown in ***Figure 4***E).

##### S2.2 Translation rate *α*

To calculate the translation rate, we simultaneously imaged the *ftz* transcription rate and the resulting Ftz protein concentration in a Ftz-MS2-LlamaTag construct (***Figure S3***A). We focused on the nuclei with no initial Ftz transcription. For these nuclei, we measured the MS2 signal (see ***Figure S3***B for a sample trace), which is an approximation of the *ftz* mRNA production rate (***Garcia et al., 2013***; ***Lammers et al., 2020***; ***Bothma et al., 2018***), and integrated this signal in order to obtain the total amount of mRNA produced (see ***Figure S3***C; ***Garcia et al. (2013***)). This integration was done by solving the differential equation for the mRNA *R*(*t*) given by

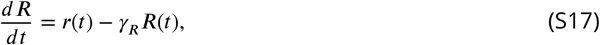

where *a*(*t*) is the transcription rate (i.e., the mRNA production rate reported by MS2 fluorescence), and *γ*_*R*_=0.099 min^−1^ (see ***Table S1***). An example of a resulting prediction for the amount of mRNA as a function of time is shown in ***Figure S3***C.

Next, we performed a parameter sweep for the translation rate *α* and, for each value of *α*, we integrated *R*(*t*) to predict the protein dynamics *P* (*t*) using the following equation (see ***Figure S3***D for sample nuclear traces)

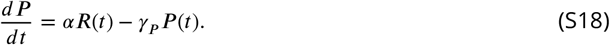

The translation rate that results in the best fit (***Figure S3***D, red line) is recorded for each nucleus. We then calculated *α* values for each embryo by averaging best-fitted *α* for each single cell (***Figure S3***E). These values are averaged across *N* = 129 cells (embryo 1) and *N* = 119 cells (embryo 2), respectively. Then we averaged the resulting value of *α* between two embryos, giving us *α* = 0.082±0.004 protein AU (mRNA AU min)^-1^ (see ***Table S1***), which is used in our dynamical systems model in the main text.

**Figure S3.**
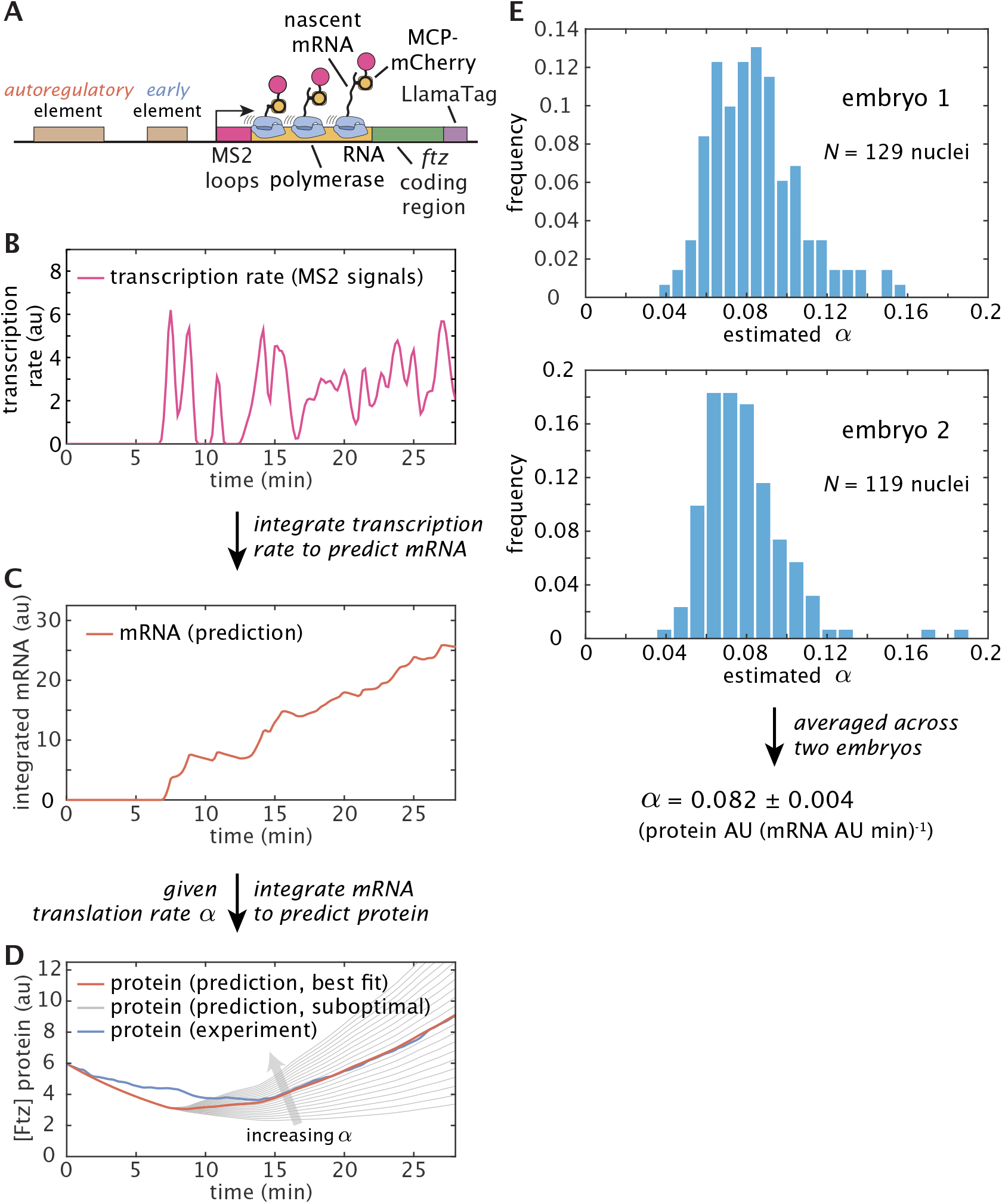
Procedure for estimating the translation rate *α*. (**A**) Ftz-MS2-LlamaTag transgenic construct used for estimating the translation rate. (**B**) Illustrative example of the raw MS2 traces that report on the instantaneous *ftz* transcription rate. MS2 traces are smoothened using a moving average of 1 min. (**C**) We integrate the MS2 traces to predict the mRNA dynamics in individual nuclei. (**D**) For a given value of the translation rate, we integrate the predicted mRNA dynamics to subsequently predict the Ftz protein dynamics in individual nuclei. The value of *α* that leads to the best agreement between prediction and experiment is found (red line). Suboptimal fits are shown as (gray lines). Ftz protein traces are smoothened using a moving average of 5 min. (**E**) Histogram distribution of best-fitted *α* values for individual cells within two embryos. Note that *α* values larger than 0.2 au are omitted for accuracy (*N* = 2 nuclei). We first calculated the average *α* for each individual embryo and then computed the average between these two embryos, resulting in *α* = 0.082 ± 0.004 protein AU (mRNA AU min)^-1^, which is used in our dynamical systems model.

**Figure S4.**
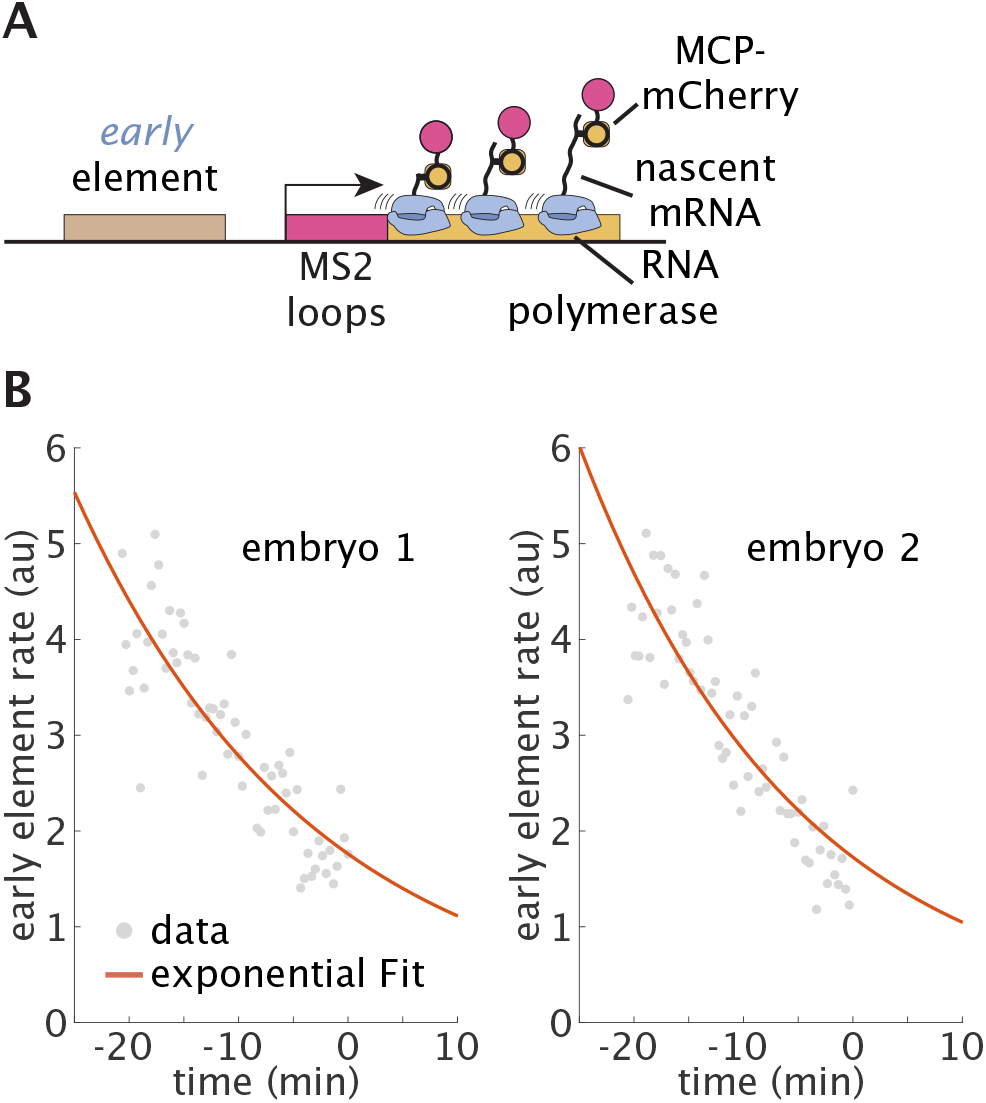
Inferring the decay rate of the early element transcriptional activity *β*. (**A**) Imaging transcriptional dynamics of the early element using the MS2 system. The construct is the same as the one described in ***Figure 3***A. (**B**) Exponential fit from two embryos. Gray dots represent the averaged early element transcription rate at individual time points from a single embryo. Red line is the exponential fit that is used to estimate the early element decay rate *β*.

##### S2.3 Early element transcriptional activity decay rate *β*

We calculated the decay in the mRNA production rate of early element, described by the decay rate *β*, from fluorescence measurements of the early element MS2 reporter construct (***Figure S4***A). We performed an exponential fit to the average trajectory of each embryo (see ***Figure S4***B for an example fit) and averaged the resulting decay rates to obtain the mean value *β* = 0.048 ± 0.0021 (see ***Table S1***, *N* = 2 embryos) that we used in the dynamical systems model.

##### S2.4 Scaling factor *c*

We estimated *c*, the ratio of maximum production rate from the endogenous locus vs. the transgene, by approximating the solution to the dynamical system in ***Eq. S4*** between two time points for which we have empirical data. In particular, we began from the experimental observation that 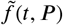 plateaus shortly before gastrulation for nuclei with total Ftz levels above about 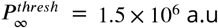.Therefore, for nuclei that satisfy 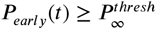 during this time (***Figure S5***A), we can approximate the nonlinear system for the autoregulatory element by the following *linear* system

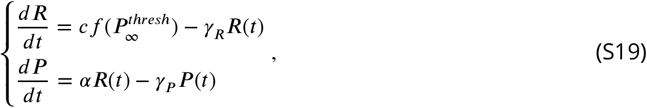

where 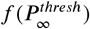is now a constant. This system of equations has an analytical solution given by

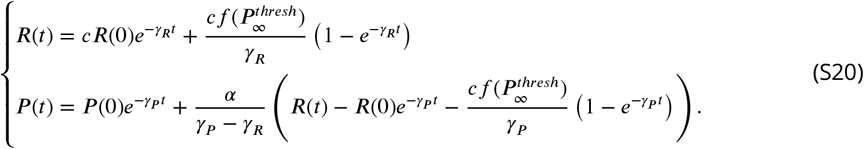

Rearranging these expressions gives

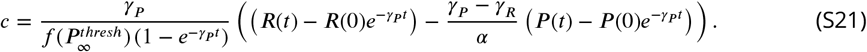

Note that here *t* = 0 is assigned to the beginning of the time window over which the simulation is performed.

We calculated the solution to ***Eq. S21*** for the individual boundary nuclei in the same embryos as used to fit the gene regulatory function (***Figure 4***D). We restricted our estimations to the 3 min before gastrulation based on personal observations that the prediction accuracy of the simulation of this linearized system tended to fall after ∼3 min. We derived the initial conditions (*R*(0), *P* (0)) from estimates of the late protein obtained by subtracting simulated early protein from the total protein trace (where the autoregulatory element was assumed to begin contributing at -20 min before gastrulation). We estimated *c* in a windowed approach whereby, for each nucleus, we simulated only over one empirical sample interval (10 s) for all intervals from 3 min before gastrulation (***Figure S5***B). We pooled all samples for individual time windows across all nuclei (***Figure S5***C and D) and averaged them to give a final estimate of *c* ≊ 0.45 ± 0.02 across the 3 min before gastrulation (where the error range is the standard error).

#### S3 Simulations

##### S3.1 Best predicted nuclei

In order to be assured of the accuracy of our conclusions concerning the dynamics of the commitment process, in ***Figure 7*** we decided to restrict our analysis to sets of nuclei whose simulated trajectories well matched the empirical traces. From our dataset, we identified such “best predicted” nuclei based on the cumulative error between a measured trajectory ({ *R*_*t*_, *P*_*t*_ }) at discrete time points *t* and a simulated trajectory 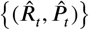 at the same time points as

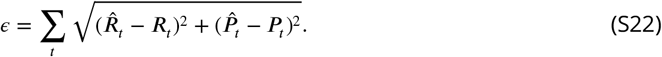

From a histogram of the errors (***Figure S6***A), which was roughly bimodal, we identified a threshold of 7 × 10^7^ to identify the 79 best predicted nuclei out of 118 nuclei total. Some sample traces from these best predicted nuclei are shown in ***Figure S6***B while traces for nuclei with high cumulative error are shown in ***Figure S6***C.

We also calculated the cumulative error when simulations were conducted for a gradual increase in autoregulatory responsiveness described by

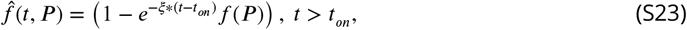

where *f* (*P*) is as defined in ***Eq. S3*** and *ξ* = 0.14 min^-1^ corresponds to a half-life of 5 min (from the observation that the autoregulatory element transitions from unresponsive to fully responsive over 10 min; see ***Figure 4***E). The resulting sample traces are shown in ***Figure S7***B. Compared to the case where we assume that the autoregulatory element becomes active instantaneously at *t*_*on*_ (***Figure S6***A), there was a slight shift in the distribution toward lower error (88 best predicted with the same cutoff as before in ***Figure S6***A), but the binary classification accuracy at gastrulation was the same. Therefore we opted to use *f* (*P*) rather than 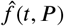 for the main analysis.

**Figure S5.**
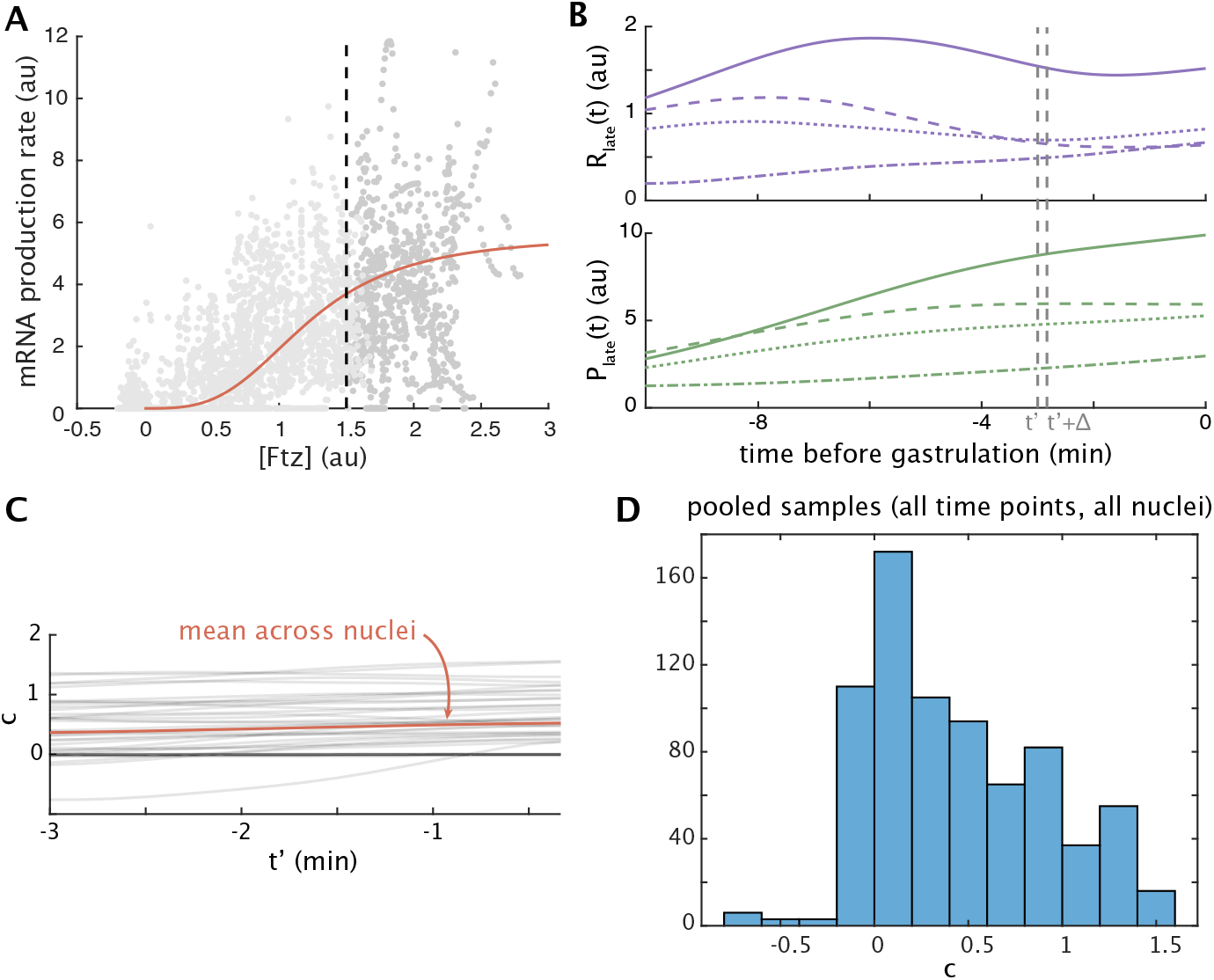
Procedure for estimating *c*, the ratio of the maximum mRNA production rate from the endogenous to the transgene. (**A**) We use a subset of samples for which the gene regulatory function is approximately saturated across the entire 3 min before gastrulation. Red, the gene regulatory function. Gray dots are individual time points from individual nuclei during the 3 min before gastrulation. Dark gray dots belong to nuclei for which 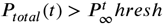 for all times between -3 and 0 min relative to gastrulation. Light gray dots correspond to nuclei that did not match this criterion. (**B**) Top, estimated late mRNA traces and bottom, estimated late protein traces for four sample nuclei drawn from the subset identified in (A). Gray dashed lines indicate one sample interval from *t*^′^ to *t*^′^ + Δ used to estimate *c* (see text). (**C**) Gray, estimates of *c* for each interval pictured in (B) and each nucleus identified in (A). Red, the mean at each time point across all nuclei. Note that most variation is between nuclei rather than across time points. (**D**) Histogram of *c* for all time intervals across all nuclei, mean 0.45, standard error 0.02.

**Figure S6.**
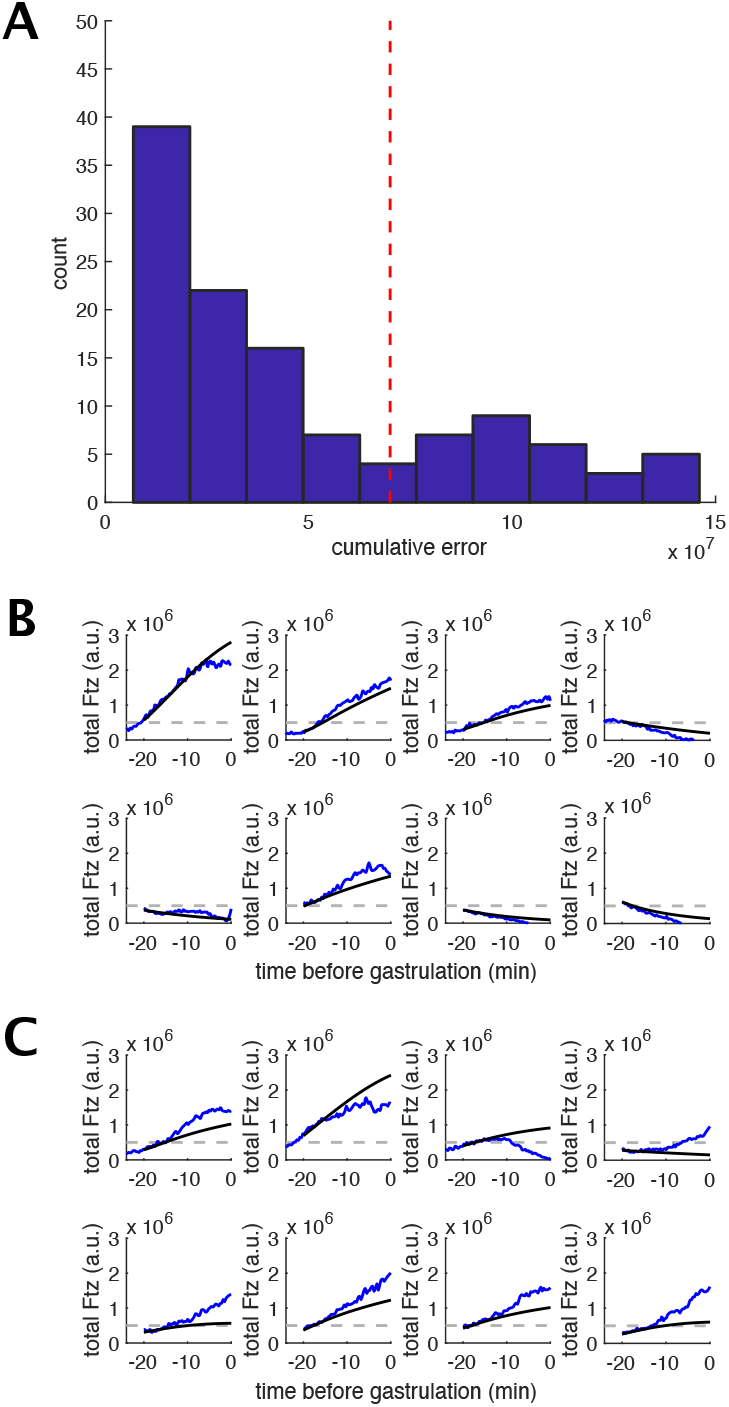
Classifying nuclei by the cumulative error in model prediction in a model of instantaneous autoregulation onset. (**A**) Histogram of cumulative errors for *N* = 118 nuclei. Simulations were conducted assuming that the autoregulatory element becomes instantaneously responsive, as in ***Eq. S3*** and the main text. (**B**) Sample traces of best predicted nuclei. In each plot, the blue curve is the measured trace and the black line is the corresponding predicted trace. The dashed gray line is the threshold for classifying a nucleus as “on” at gastrulation. (**C**) As for (B), but for nuclei that did not qualify to be best predicted.

**Figure S7.**
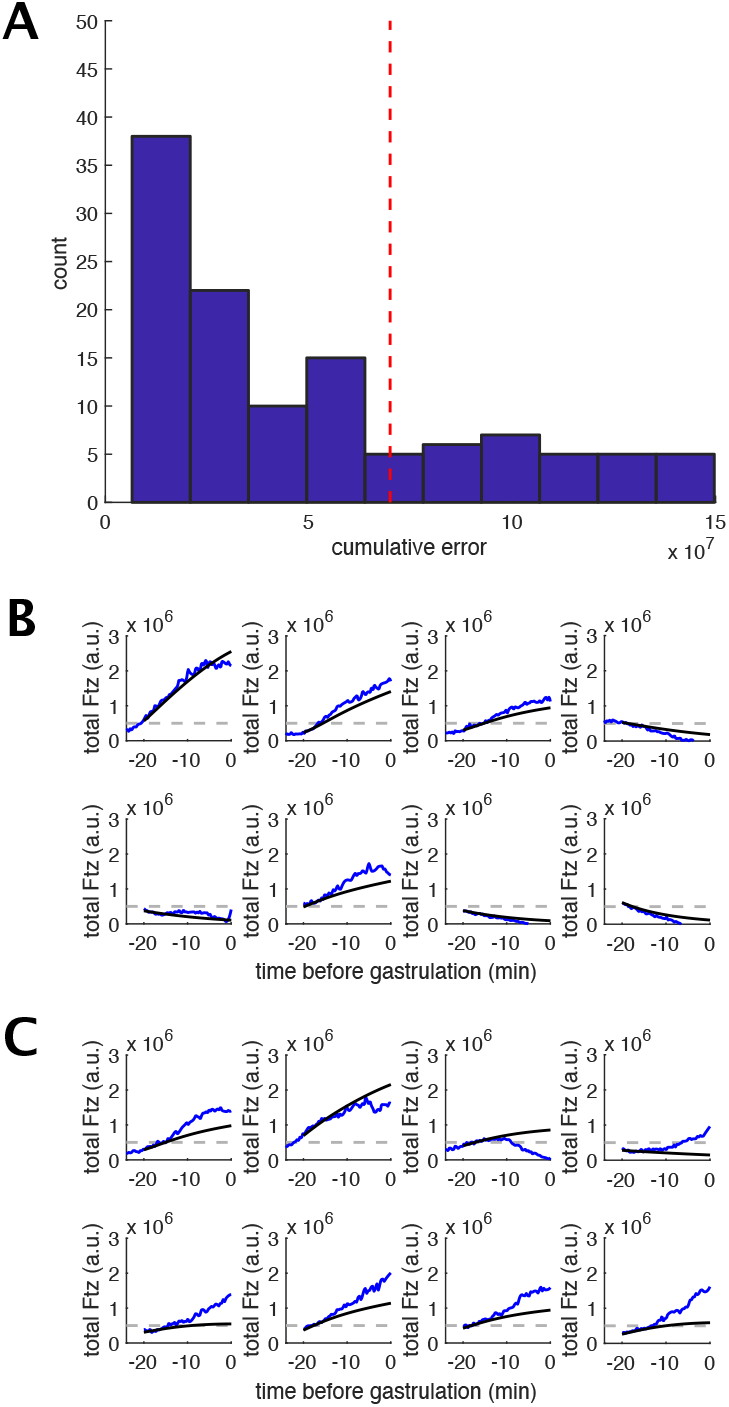
Classifying nuclei by the cumulative error in model prediction in a model of gradual autoregulation onset. (**A**) Histogram of cumulative errors for *N* = 118 nuclei. Simulations were conducted assuming a gradual increase in responsiveness of the autoregulatory element, following the dynamics in ***Eq. S23***. (**B**) Sample traces of best predicted nuclei. In each plot, the blue curve is the measured trace and the black line is the corresponding predicted trace. The dashed gray line is the threshold for classifying a nucleus as “on” at gastrulation. (**C**) As for (B), but for nuclei that did not qualify to be best predicted.

**Figure S8.**
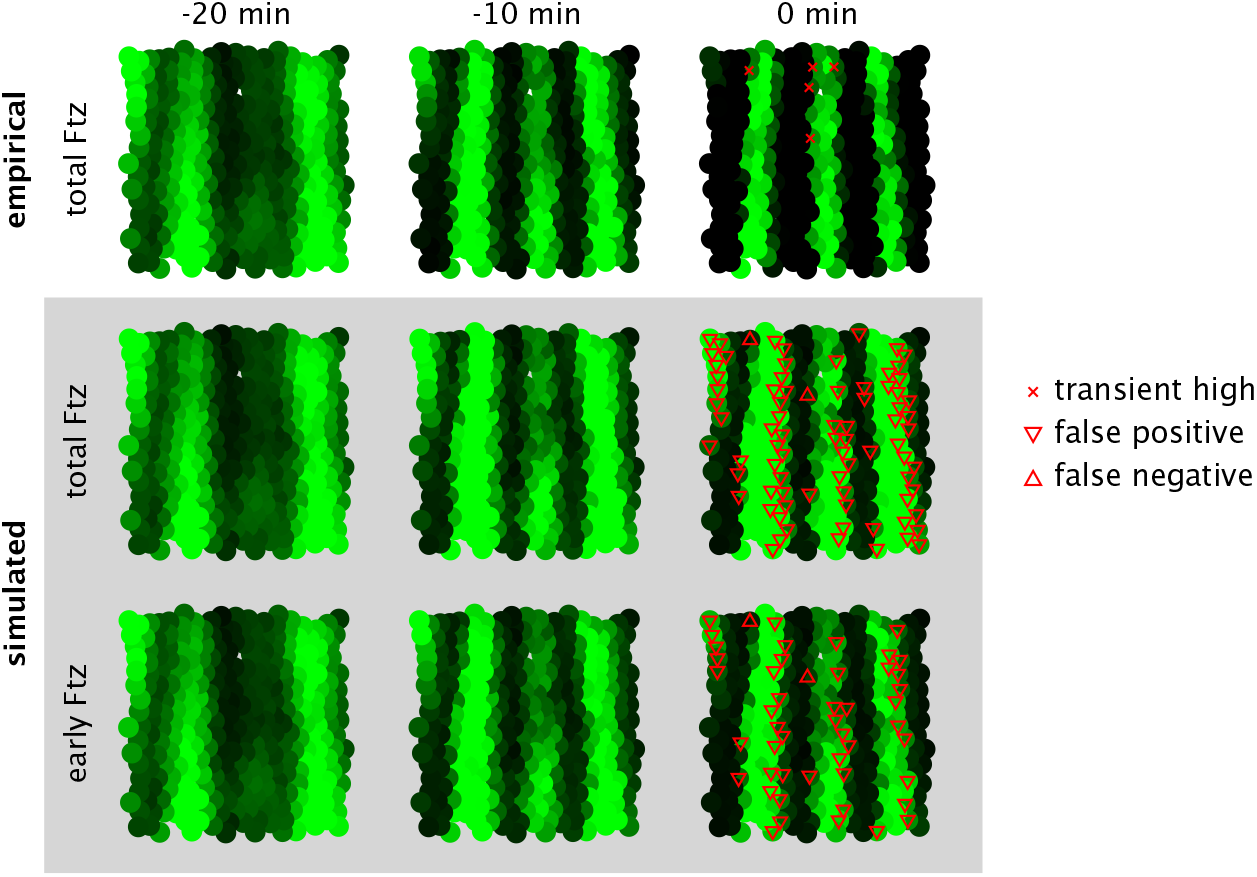
Whole-embryo simulation for the Ftz stripe pattern. Illustrative whole-embryo simulation for the same embryo as pictured in ***Figure 6***C. Pixel intensities are normalized to the steady-state high Ftz level.

##### S3.2 Whole-embryo simulations

We were curious about how accurately our model, which is based on measurements at the anterior boundary of stripe 4, would predict Ftz state for all nuclei spanning stripes 3, 4 and 5 measured during our experiments. Assuming all nuclei follow the same dynamics as given in ***Eq. 5*** and ***Eq. S4***, we repeated the analysis from ***Figure 5*** to predict Ftz concentration at gastrulation. We achieved a binary classification accuracy of (74.6%, or 763 of 1036 nuclei). Interestingly, this is worse than the accuracy achieved from thresholding early protein alone (84.3%, or 873 of 1036 nuclei), which is itself comparably accurate to the predictions for the anterior boundary of stripe 4 (with or without the autoregulatory contribution). The bulk of classification errors for the whole-embryo simulations, whether from thresholding full simulations or thresholding early protein alone, were false positives at the posterior boundaries of stripes, as in the example plotted in ***Figure S8***. This indicates that something differs in the regulation of Ftz at the anterior boundaries of stripes as compared to the posterior boundaries. For example, it has been noted that the posterior, but not the anterior, boundaries of Ftz stripes are repressed by *sloppy paired* (*slp*) (***Clark, 2017***).

##### S3.3 Delaying the onset of responsiveness of the autoregulatory element

From a mathematical standpoint, we can treat a delay in *t*_*on*_ as a change in the starting time of the simulation, which introduces a corresponding change to the initial conditions (*r*(*t*_*on*_), *R*_*early*_(*t*_*on*_), *P*_*early*_(*t*_*on*_)) of the early module. In this way, the time at which the trajectory of the early module (*r*(*t*), *R*_*early*_(*t*), *P*_*early*_(*t*)) crosses the switching separatrix is the latest time at which the autoregulatory element can become responsive and still commit a cell to the appropriate (high) fate (***Figure S9***). This follows from three conditions: (1) the early element is time invariant, (2) the early element is independent of the autoregulatory module (the same is not true for the autoregulatory module, which takes the output of the early module as its input), and (3) Ftz fate corresponds to the Ftz concentration state at infinite time. (1) and (2) ensure that, even if we delay the autoregulatory element, we can continue to use ***Eq. 5*** to simulate early protein by just changing the initial conditions, while (3) ensures that the delayed start of the autoregulatory element will not change the location of the switching surface (which is relative to steady state, not to a transient state of the trajectory at a fixed point in time).

**Figure S9.**
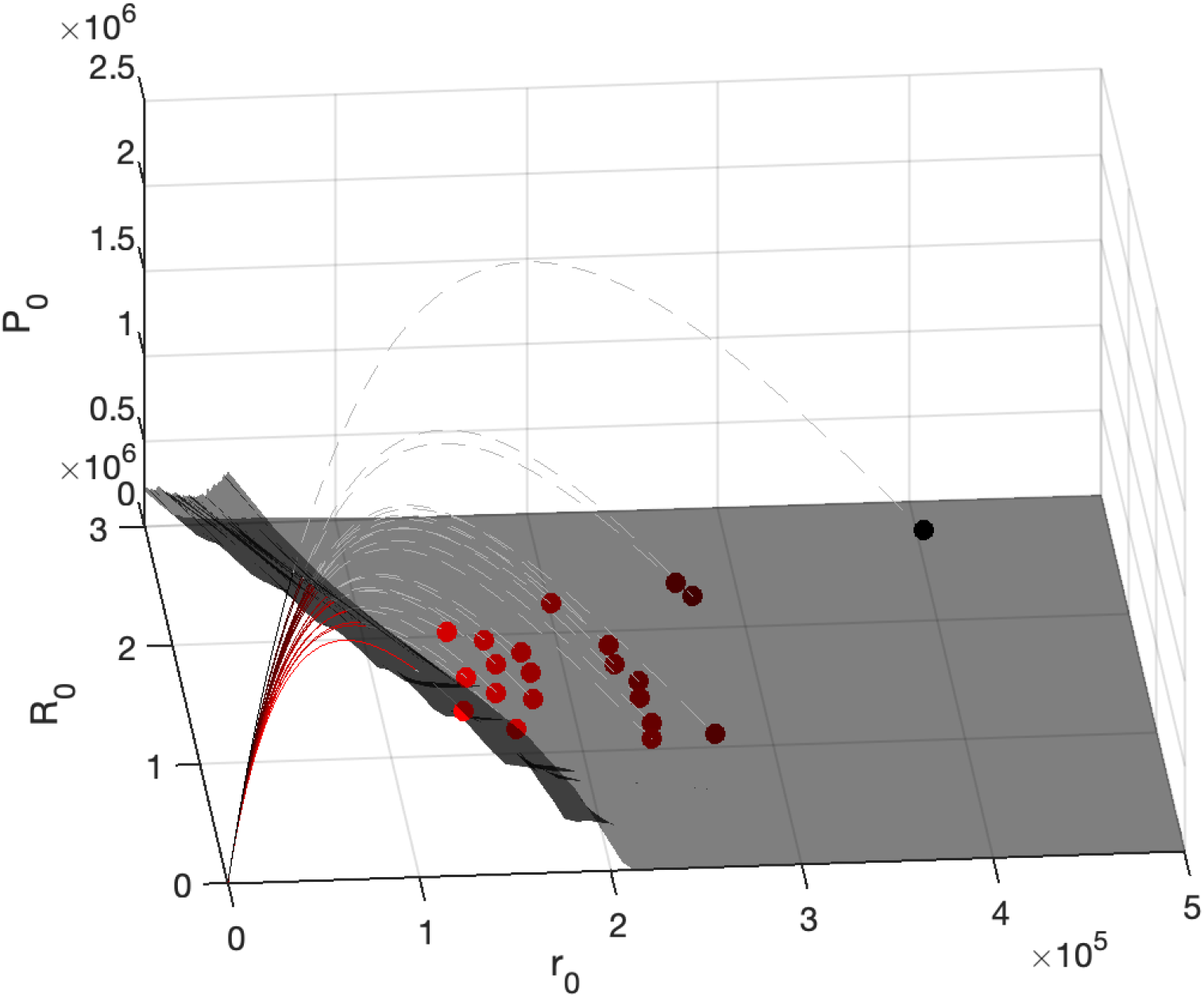
Sensitivity of Ftz fate to the autoregulatory onset time *t*_*on*_. Dashed gray lines denote the trajectories (*r*(*t*),*R*_*early*_(*t*),*P*_*early*_(*t*)) corresponding to each nucleus. Trajectories begin at the colored points, each of which corresponds to the state of the early module in a high-fated nucleus (*N* = 21 same as analyzed in ***Figure 7***) at time -20 min. Nuclei are colored based on the amount of time until the trajectory crosses the switching separatrix from ***Figure 6***B, with darker colors indicating longer times. As explained in the text, this crossing time corresponds to the longest acceptable delay in the onset of autoregulatory responsiveness without changing the fate of the cell. Intriguingly, the trajectories of early protein run parallel to the separatrix before converging to cross it in a restricted region of parameter space.

In ***Figure 7*** we analyze the commitment window by varying *t*_*off*_ at the same time as *t*_*on*_. In ***Figure S10*** we report full results for simultaneous variation in *t*_*on*_ and *t*_*off*_. The strictness with which cell fate must be specified determines the variation in timing that can be tolerated. For example, if the early element ceases production at or after gastrulation (*t*_*off*_ > 0), the autoregulatory element can delay responsiveness until -15 min and still guide at least 75% of cells to the appropriate fate. If the early element does not cease production until 20 min, then the autoregulatory element may turn on just after -6 min and still direct 75% of cells to the correct fate. From these results, we see that almost no cells commit to the high fate when *t*_*on*_ > *t*_*off*_. For this reason, in the main text we always set *t*_*off*_ > *t*_*on*_.

##### S3.4 Stochastic simulations

Stochasticity in gene expression during embryonic development can compromise or improve system function depending on the context (***Zhang et al., 2012***; ***Papadopoulos et al., 2019***). We sought to investigate whether stochasticity in gene expression (1) was sufficient to explain the prediction error rates of our deterministic models, and (2) could drive stochastic switching at appreciable rates. We examined these questions using stochastic differential equations (SDEs), assuming that the noise in our data arises solely from stochastic dynamics within the cells rather than from measurement noise. Generally, we expect this method to overestimate the error.

**Figure S10.**
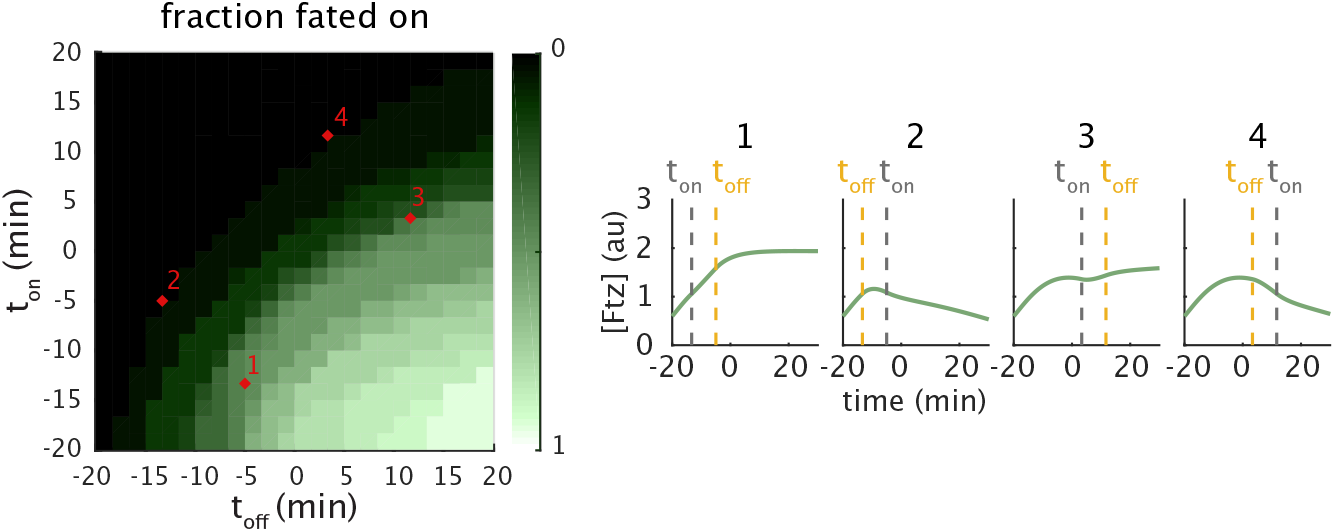
Dependence of Ftz fate adoption on the time that autoregulatory responsiveness begins and production rate from the early element shuts down. Left, heat map showing the fraction of cells predicted to adopt the high Ftz fate for different combinations of *t*_*on*_ and *t*_*off*_. For this fraction to achieve 95% requires fairly strict coordination between the timing of modules, with neither event deviating more than about 7 min from *t*_*on*_ = −20 min or *t*_*off*_ = 20 min (on the order of the mRNA decay rate). Allowing for larger percentages of errors in fate determination significantly weakens constraints on regulatory timing, with a variation of up to 15 min in *t*_*off*_ or *t*_*on*_ still resulting in roughly 50% of cells adopting the appropriate high fate. Right, examples of simulated Ftz traces for a single nucleus with the timing conditions corresponding to the red marks on the heatmap.

In a stochastic differential equation model, changes in the amount of late RNA *dR*_*late*_ and total protein *dP*_*total*_ over a time interval *dt* are given by

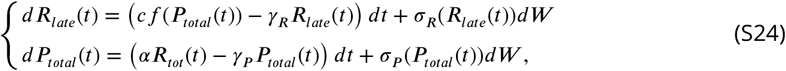

where *dW* is Gaussian with mean 0 and variance *dt* (***Gillespie, 2007***). The terms *σ*_*R*_(*R*_*late*_) and *σ*_*P*_ (*P*_*late*_) scale the variance of the noise from one time increment to the next.

For this analysis, we used our experimental setup featuring endogenous Ftz-LlamaTag driving an autoregulatory element transgene tagged with MS2 as introduced in ***Figure 4***. Because, over a small time interval where mRNA degradation is negligible, the MS2 signal reports on the rate of mRNA production, this signal gave us direct access to *dR*_*late*_(*t*_*i*_) at discrete time points *t*_*i*_. As a result, we can use this measure of *dR*_*late*_(*t*) to estimate for the late mRNA *R*_*late*_(*t*) by integrating the MS2 signal following

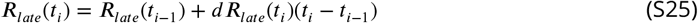

under the assumption that *R*_*late*_(*t*_*on*_) = 0 (meaning that the autoregulatory element only becomes responsive at time *t*_*on*_). Since we also have direct measurements of *P*_*total*_(*t*), this allows us to rearrange ***Eq. S24*** so as to estimate the noise contribution *σ*_*R*_(*R*_*late*_(*t*))*dW* from

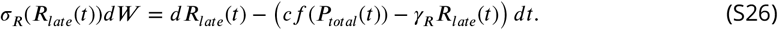

If we further assume that *R*_*early*_(*t*) is noiseless and therefore given by the deterministic solution in ***Eq. S9***, we can estimate the total mRNA as *R*_*total*_(*t*) = *R*_*early*_(*t*) + *R*_*late*_(*t*). Then, since we have simultaneous measurements of total Ftz protein *P*_*total*_, we can also rearrange the lower equation in ***Eq. S24*** to estimate *σ*_*P*_ (*P*_*total*_(*t*))*dW* from

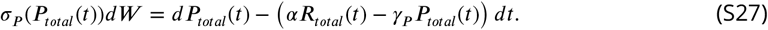

**Figure S11.**
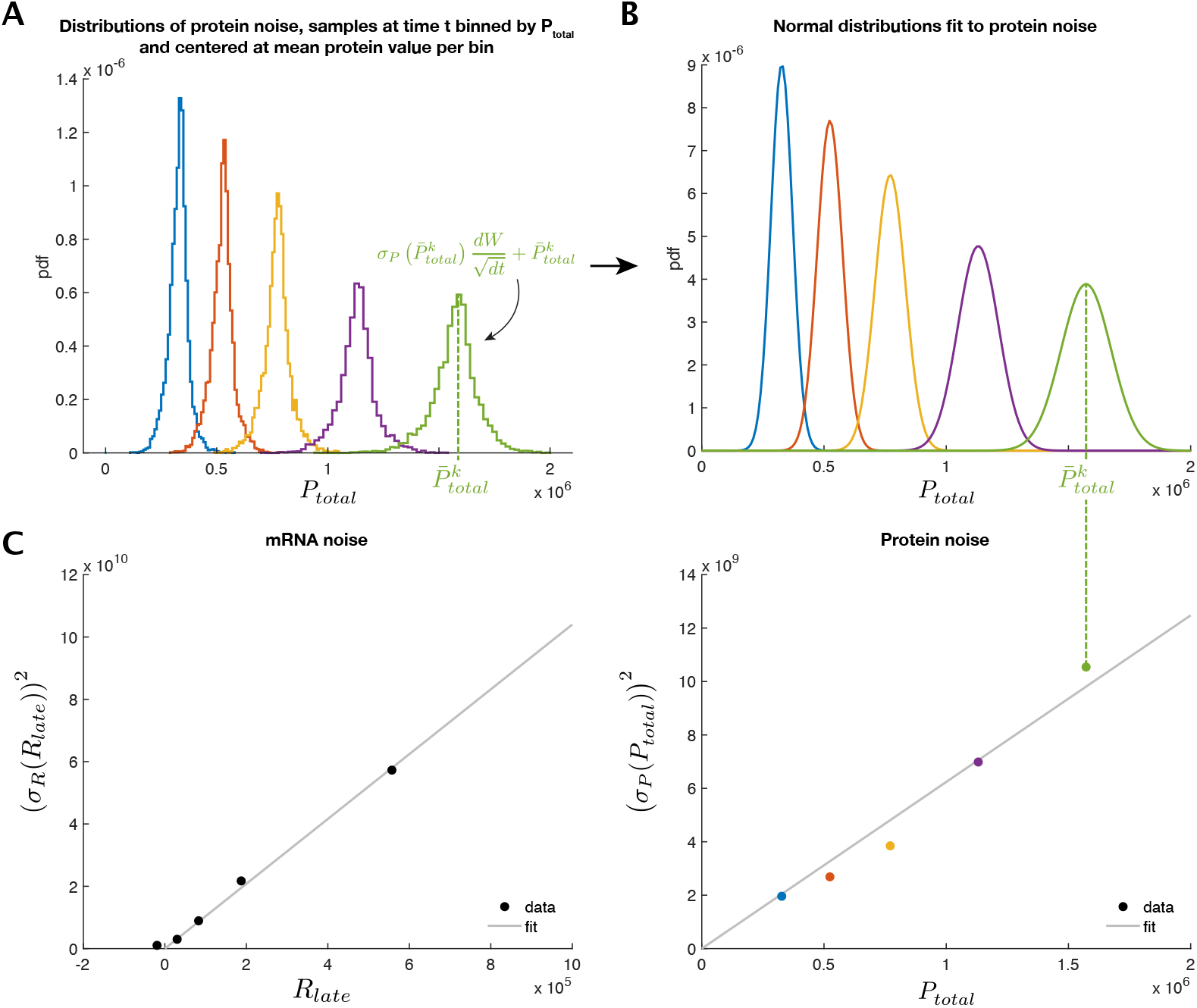
Procedure for estimating mRNA and protein expression noise. (**A**) We extract samples of the noise contribution for an individual sample point *σ*_*P*_ (*P*_*total*_)*dW* following the method described in the text. We bin the samples by the corresponding *P*_*total*_ and calculate the mean protein level 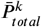 within each protein concentration bin *k*. We divide the samples by 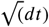 to remove the variance contribution from the *dW* term in ***Eq. S24***. We centered the samples from each bin at the corresponding 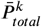 to generate the pictured histograms. (**B**) A normal distribution is fitted to each bin in panel (A). (**C**) Right, we estimate *σ*_*P*_ (*P*) by fitting a line to the relationship between 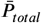 and variance of the distribution fitted to the corresponding bin in (A) and (B). Left, the equivalent result for mRNA noise, which is calculated identically except that results are binned by *R*_*late*_(*t*) (not pictured).

We can perform the above analysis on individual measured traces to produce a large number of sample points of *σ*_*R*_(*R*_*late*_)*dW* and *σ*_*P*_ (*P*_*total*_)*dW*. With these data we will aim to estimate *σ*_*R*_(*R*) and *σ*_*P*_ (*P*). We assume the noise characteristics are time invariant, which allows us to pool all samples at all time points and bin them by the corresponding *R*_*late*_ or *P*_*total*_. For example, for protein, we treat each protein concentration bin *k* as a population of samples of 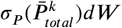where 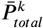 is the mean of the samples in bin *k* (***Figure S11***A). Since we assume *dW* is normally distributed with variance *dt*, we divide all sample values by 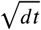 and fit a normal distribution to the resulting distribution within each protein concentration bin (***Figure S11***B). We found that the variances *σ*_*P*_ (*P*)^2^ are quite well approximated by a linear relation *σ*_*P*_ (*P*)^2^ = *aP* + *b* (***Figure S11***C, right). The mRNA variance was estimated similarly and also found to fit a linear relation (***Figure S11***C, left). Noise was estimated from all available trajectories, regardless of whether they were part of the stripe 4 anterior boundary.

Having estimated the noise, we investigated whether stochasticity could explain the error rate in our predictions of Ftz expression state at gastrulation. We simulated *N*_*sim*_ = 100 experiments, each consisting of *N*_*nuc*_ = 118 nuclei evenly split between those deterministically predicted to be on and those deterministically predicted to be off. Specifically, we found the convex hull defined by the experimentally measured initial conditions for nuclei at the stripe 4 anterior boundary, and drew random initial conditions for the stochastically simulated nuclei from a uniform distribution within this hull. We assigned to each simulated experiment 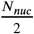 points below the blue surface in ***Figure 5***C and 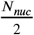 above the surface without replacement (i.e., every nucleus in every simulated experiment has a unique set of initial conditions). Individual stochastic trajectories were generated using the Euler-Maruyama method, with the modification that protein and mRNA concentrations were forcibly lower bounded at 0 (i.e., random fluctuations that would bring concentrations to negative values were capped to instead bring the concentration to zero). Trajectories were simulated for 20 min until gastrulation and thresholded with the same value as for our deterministic simulations (***Figure S12***).

**Figure S12.**
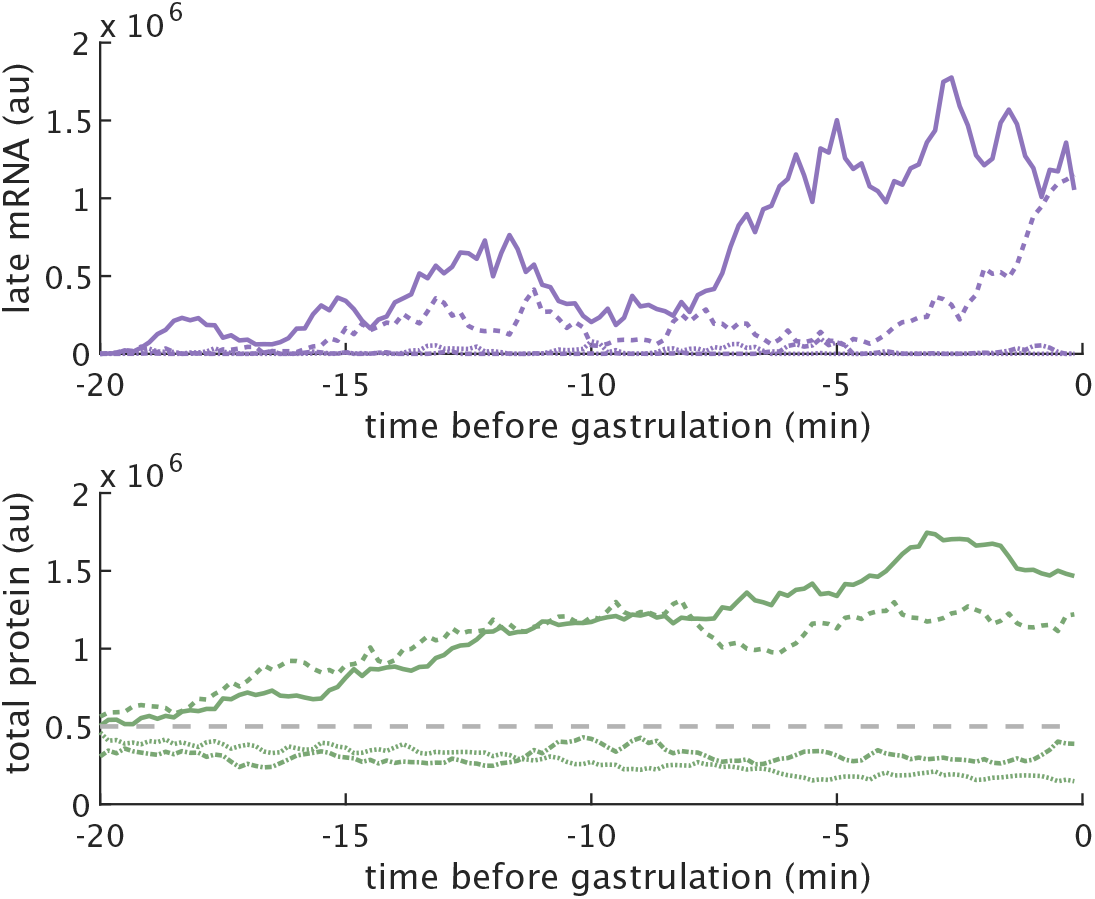
Sample traces for nuclei simulated with stochastic dynamics. Top, late mRNA and bottom, total protein traces simulated for individual nuclei obeying the stochastic dynamics in ***Eq. S24***. Gray dashed line in the bottom plot is the threshold to be classified as high Ftz at gastrulation.

From our *N*_*sim*_ = 100 simulated experiments, we calculated a distribution of error rates given by

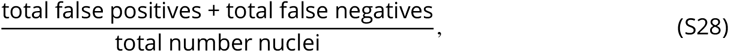

where we compare the predicted outcomes from deterministic simulations to the “ground truth” of the stochastic simulations.

In ***Figure S13*** we plot the cumulative distribution of error rates for our simulations (black), broken down into false negatives (red) and false positives (green). The dashed vertical lines indicate the experimentally measured error rates with the same color code. Where the vertical lines intersect the corresponding cumulative distributions indicates the probability of measuring an error rate up to that rate. If the system is really described by the stochastic dynamics we have inferred, then the most likely error rates are those that intersect the curves where their slope is highest.

We found that the empirical error rate across 3 embryos was roughly twice that of the most likely error rates from our stochastic simulations corresponding to the middle of calculated cumulative distribution functions (***Figure S13***, left), with the majority of errors being false negatives. We knew from observation that one embryo had a large number of false negative predictions (***Section 2.3***, and, interestingly, if we exclude this embryo from analysis, then the empirical error rate aligns well with what the stochastic model predicts to be most likely (***Figure S13***, right). This result suggests that many of our prediction errors can likely be attributed to stochastic fluctuations.

**Figure S13.**
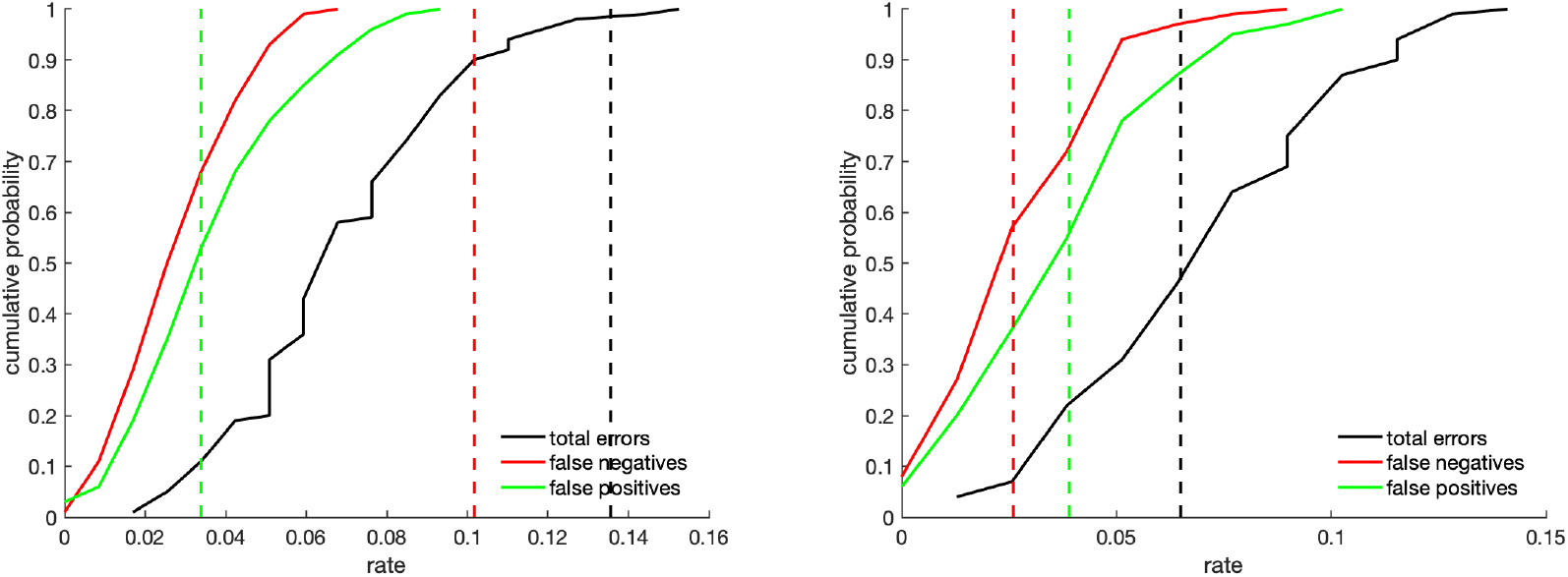
Error rates resulting from deterministic predictions made when gene expression is stochastic. Solid lines, cumulative distribution functions for error rates in the prediction of results for *N* = 100 simulated stochastic experiments (using the stochastic differential equation in ***Eq. S24***), when using a deterministic simulation to make the predictions. For each simulation, we predicted the trajectories of *N*_*nuc*_ boundary nuclei evenly partitioned between starting states above and below the surface for classifying high or low Ftz at gastrulation (***Figure 5***). Dotted vertical lines correspond to empirical error rates for three embryos with *N*_*nuc*_ = 118 nuclei (left) across 3 embryos, or excluding one embryo that had a high number of deterministic false negatives (***Section 2.3***) for *N*_*nuc*_ = 78 nuclei (right) across 2 embryos.

Having determined that the error of the model in predicting Ftz expression state at gastrulation is comparable to the error expected when considering gene expression stochasticity, we next turned to the question of whether gene expression stochasticity is expected to play a large role in the long-term Ftz fate of cells in which the early element is no longer active. We ran *N* = 100 simulations beginning from the high steady state and calculated the distribution of first-passage times to particular protein values (***Figure S14***, left) or to within some Euclidean distance of the opposite steady state (***Figure S14***, right). These measures give an approximation of the switching rate depending upon how stringently one defines a threshold for switching.

Both trends indicate that switching from high to low occurs at a much faster rate than low to high, with conservative rates of stochastic switching between the high and low Ftz fates of around 2 hr and between low and high fates of approximately 3 hr. For comparison of timescales, Ftz stripes are no longer experimentally detected before the end of germband extension (***Hafen et al., 1984***; ***Carroll, 1985***), which occurs approximately 1.5 hours after gastrulation (***da Silva and Vincent, 2007***). Thus, we find no strong evidence that stochastic switching should contribute significantly to Ftz stripe patterning.

**Figure S14.**
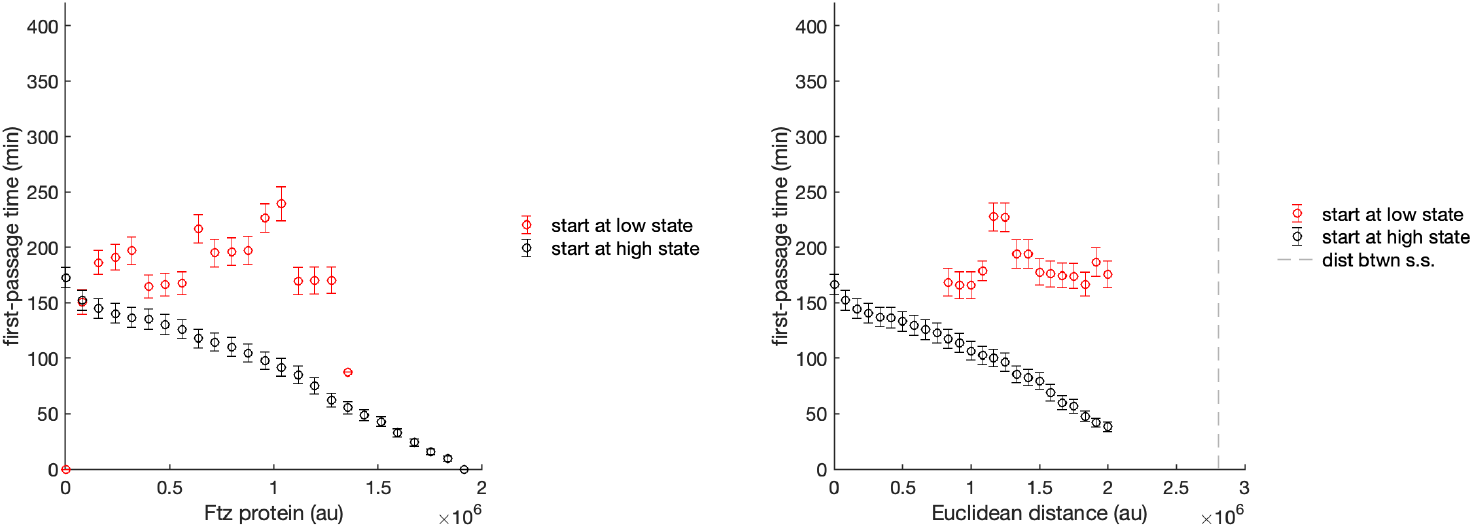
First-passage times suggest that the stochastic switching rate between Ftz fates at steady state occurs on a timescale of hours. Left, time to first passage of a stochastic trace starting from the high (black) or low (red) fate to a given protein value. Right, time to first passage of a stochastic trajectory with a given Euclidean distance from the opposite steady state. In both plots, red denotes simulations beginning in the low Ftz state and black denotes simulations beginning in the high Ftz state.

## References

Alon, U. (2006). An Introduction to Systems Biology. Chapman and Hall/CRC.

Alon, U. (2007). Network motifs: Theory and experimental approaches. Nature Reviews Genetics, 8:450–461.

Angeli, D., Ferrell, J. E., and Sontag, E. D. (2004). Detection of multistability, bifurcations, and hysteresis in a large class of biological positive-feedback systems. Proceedings of the National Academy of Sciences, 101(7):1822– 1827.

Bending, D., Martín, P. P., Paduraru, A., Ducker, C., Marzaganov, E., Laviron, M., Kitano, S., Miyachi, H., Crompton, T., and Ono, M. (2018). A timer for analyzing temporally dynamic changes in transcription during differentiation in vivo. Journal of Cell Biology, 217(8):2931–2950.

Blanchini, F. and Franco, E. (2014). Structural analysis of biological networks. In A Systems Theoretic Approach to Systems and Synthetic Biology I: Models and System Characterizations, pages 47–71. Springer.

Bolouri, H. and Davidson, E. H. (2002). Modeling transcriptional regulatory networks. BioEssays, 24(12):1118– 1129.

Bothma, J. P., Garcia, H. G., Esposito, E., Schlissel, G., Gregor, T., and Levine, M. (2014). Dynamic regulation of eve stripe 2 expression reveals transcriptional bursts in living Drosophila embryos. Proceedings of the National Academy of Sciences, 111(29):10598–10603.

Bothma, J. P., Norstad, M. R., Alamos, S., and Garcia, H. G. (2018). LlamaTags: A versatile tool to image transcription factor dynamics in live embryos. Cell, 173(7):1810–1822.

Bouchoucha, Y. X., Reingruber, J., Labalette, C., Wassef, M. A., Thierion, E., Desmarquet-Trin Dinh, C., Holcman, D., Gilardi-Hebenstreit, P., and Charnay, P. (2013). Dissection of a Krox20 positive feedback loop driving cell fate choices in hindbrain patterning. Molecular Systems Biology, 9:690.

Carroll, S. (1985). Localization of the fushi tarazu protein during Drosophila embryogenesis. Cell, 43(1):47–57.

Clark, E. (2017). Dynamic patterning by the Drosophila pair-rule network reconciles long-germ and short-germ segmentation. PLoS biology, 15(9):e2002439.

Clark, E. and Akam, M. (2016). Odd-paired controls frequency doubling in Drosophila segmentation by altering the pair-rule gene regulatory network. eLife, 5:e18215.

Clark, E., Battistara, M., and Benton, M. A. (2022). A timer gene network is spatially regulated by the terminal system in the Drosophila embryo. bioRxiv.

Clark, E. and Peel, A. D. (2018). Evidence for the temporal regulation of insect segmentation by a conserved sequence of transcription factors. Development, 145(10).

Corson, F. and Siggia, E. D. (2017). Gene-free methodology for cell fate dynamics during development. eLife, 6:e30743.

Cosentino, C. and Bates, D. (2011). Feedback Control in Systems Biology. CRC Press.

Cotterell, J. and Sharpe, J. (2010). An atlas of gene regulatory networks reveals multiple three-gene mechanisms for interpreting morphogen gradients. Molecular Systems Biology, 6:425.

Crews, S. T. and Pearson, J. C. (2009). Transcriptional autoregulation in development. Current Biology, 19(6):R241–R246.

da Silva, S. M. and Vincent, J.-P. (2007). Oriented cell divisions in the extending germband of Drosophila. Development, 134(17):3049–3054.

Dalton, S. (2015). Linking the cell cycle to cell fate decisions. Trends in Cell Biology, 25(10):592–600.

Dearolf, C. R., Topol, J., and Parker, C. S. (1989). Transcriptional control of Drosophila fushi tarazu zebra stripe expression. Genes & development, 3(3):384–398.

Del Vecchio, D., Densmore, D., El-Samad, H., Ingber, D., Khalil, A. S., Kosuri Sri Lander, A. D., and Tang, C. (2016). What have the principles of engineering taught us about biological systems? Cell Systems, 2:5–7.

Dubuis, J. O., Tkačik, G., Wieschaus, E. F., Gregor, T., and Bialek, W. (2013). Positional information, in bits. Proceedings of the National Academy of Sciences, 110(41):16301–16308.

Edgar, B. A., Weir, M. P., Schubiger, G., and Kornberg, T. (1986). Repression and turnover pattern fushi tarazu RNA in the early Drosophila embryo. Cell, 47(5):747–754.

Ferrell, J. E. (2002). Self-perpetuating states in signal transduction: positive feedback, double-negative feedback and bistability. Current Opinion in Cell Biology, 14(2):140–148.

Furusawa, C. and Kaneko, K. (2012). A dynamical-systems view of stem cell biology. Science, 338(6104):215–217.

Garcia, H. G., Tikhonov, M., Lin, A., and Gregor, T. (2013). Quantitative imaging of transcription in living Drosophila embryos links polymerase activity to patterning. Current Biology, 23(21):2140–2145.

Gillespie, D. T. (2007). Stochastic simulation of chemical kinetics. Annual Review of Physical Chemistry, 58(1):35– 55.

Goglia, A. G. and Toettcher, J. E. (2019). A bright future: Optogenetics to dissect the spatiotemporal control of cell behavior. Current Opinion in Chemical Biology, 48:106–113.

Graham, P. L., Fischer, M. D., Giri, A., and Pick, L. (2021). The fushi tarazu zebra element is not required for Drosophila viability or fertility. G3, 11(11).

Graham, T. G. W., Tabei, S. M. A., Dinner, A. R., and Rebay, I. (2010). Modeling bistable cell-fate choices in the Drosophila eye: qualitative and quantitative perspectives. Development, 137(14):2265–2278.

Gratz, S. J., Rubinstein, C. D., Harrison, M. M., Wildonger, J., and O’Connor-Giles, K. M. (2015). CRISPR-Cas9 genome editing in Drosophila. Current Protocols in Molecular Biology, 111(1):1–31.

Gregor, T., Tank, D. W., Wieschaus, E. F., and Bialek, W. (2007). Probing the limits to positional information. Cell, 130(1):153–164.

Gutenkunst, R. N., Waterfall, J. J., Casey, F. P., Brown, K. S., Myers, C. R., and Sethna, J. P. (2007). Universally sloppy parameter sensitivities in systems biology models. PLoS Computational Biology, 3(10):e189.

Hafen, E., Kuroiwa, A., and Gehring, W. J. (1984). Spatial distribution of transcripts from the segmentation gene fushi tarazu during Drosophila embryonic development. Cell, 37:833–841.

Hartwell, L. H., Hopfield, J. J., Leibler, S., and Murray, A. W. (1999). From molecular to modular cell biology. Nature, 402:C47–C52.

Hiromi, Y. and Gehring, W. J. (1987). Regulation and function of the Drosophila segmentation gene fushi tarazu. Cell, 50(6):963–974.

Hiromi, Y., Kuroiwa, A., and Gehring, W. J. (1985). Control elements of the Drosophila segmentation gene fushi tarazu. Cell, 43(3):603–613.

Ingram, P. J., Stumpf, M. P. H., and Stark, J. (2006). Network motifs: structure does not determine function. BMC Genomics, 7:108.

Jaeger, J. and Monk, N. (2014). Bioattractors: Dynamical systems theory and the evolution of regulatory processes. The Journal of Physiology, 592(11):2267–2281.

Jaeger, J., Surkova, S., Blagov, M., Janssens, H., Kosman, D., Kozlov, K. N., Manu Myasnikova, E., Vanario-Alonso, C. E., Samsonova, M., Sharp, D. H., and Reinitz, J. (2004). Dynamic control of positional information in the early Drosophila embryo. Nature, 430:368–371.

Johnson, H. E. and Toettcher, J. E. (2018). Illuminating developmental biology with cellular optogenetics. Current Opinion in Biotechnology, 52:42–48.

Khammash, M. (2016). An engineering viewpoint on biological robustness. BMC Biology, 14:22.

Kim, Y. J., Rhee, K., Liu, J., Jeammet, P., Turner, M., Small, S., and Garcia, H. G. (2021). Predictive modeling reveals that higher-order cooperativity drives transcriptional repression in a synthetic developmental enhancer. bioRxiv.

Kueh, H. Y., Champhekar, A., Nutt, S. L., Elowitz, M. B., and Rothenberg, E. V. (2013). Positive feedback between PU.1 and the cell cycle controls myeloid differentiation. Science, 341(6146):670–673.

Lai, K., Robertson, M. J., and Schaffer, D. V. (2004). The Sonic Hedgehog signaling system as a bistable genetic switch. Biophysical Journal, 86(5):2748–2757.

Lammers, N. C., Galstyan, V., Reimer, A., Medin, S. A., Wiggins, C. H., and Garcia, H. G. (2020). Multimodal transcriptional control of pattern formation in embryonic development. Proceedings of the National Academy of Sciences of the United States of America, 117(2):836–847.

Laslo, P., Spooner, C. J., Warmflash, A., Lancki, D. W., Lee, H.-J., Sciammas, R., Gantner, B. N., Dinner, A. R., and Singh, H. (2006). Multilineage transcriptional priming and determination of alternate hematopoietic cell fates. Cell, 126(4):755–766.

Lopes, F. J. P., Vieira, F. M. C., Holloway, D. M., Bisch, P. M., and Spirov, A. V. (2008). Spatial bistability generates hunchback expression sharpness in the Drosophila embryo. PLoS Computational Biology, 4(9):e1000184.

Lucas, T., Ferraro, T., Roelens, B., De Las Heras Chanes, J., Walczak, A. M., Coppey, M., and Dostatni, N. (2013). Live imaging of Bicoid-dependent transcription in Drosophila embryos. Current Biology, 23(21):2135–2139.

MacArthur, S., Li, X.-Y., Li, J., Brown, J. B., Chu, H. C., Zeng, L., Grondona, B. P., Hechmer, A., Simirenko, L., Keränen, S. V., Knowles, D. W., Stapleton, M., Bickel, P., Biggin, M. D., and Eisen, M. B. (2009). Developmental roles of 21 Drosophila transcription factors are determined by quantitative differences in binding to an overlapping set of thousands of genomic regions. Genome Biology, 10(7):R80.

Manu Surkova, S., Spirov, A. V., Gursky, V. V., Janssens, H., Kim, A.-R., Radulescu, O., Vanario-Alonso, C. E., Sharp, D. H., Samsonova, M., and Reinitz, J. (2009). Canalization of gene expression in the Drosophila blastoderm by gap gene cross regulation. PLoS Biology, 7(3):e1000049.

McNamara, H. M., Ramm, B., and Toettcher, J. E. (2022). Synthetic developmental biology: New tools to deconstruct and rebuild developmental systems. Seminars in Cell & Developmental Biology.

McNeely, K. C. and Dwyer, N. D. (2021). Cytokinetic abscission regulation in neural stem cells and tissue development. Current Stem Cell Reports, 7(4):161–173.

Nüsslein-Volhard, C. and Wieschaus, E. (1980). Mutations affecting segment number and polarity in Drosophila. Nature, 287(5785):795–801.

Papadopoulos, D. K., Skouloudaki, K., Engström, Y., Terenius, L., Rigler, R., Zechner, C., Vukojević, V., and Tomancak, P. (2019). Control of Hox transcription factor concentration and cell-to-cell variability by an autoregulatory switch. Development.

Papatsenko, D. and Levine, M. (2011). The Drosophila gap gene network is composed of two parallel toggle switches. PLoS ONE, 6(7):e21145.

Peter, I. S. and Davidson, E. H. (2015). Genomic control process: development and evolution. Academic Press.

Petkova, M. D., Tkačik, G., Bialek, W., Wieschaus, E. F., and Gregor, T. (2019). Optimal decoding of cellular identities in a genetic network. Cell, 176(4):844–855.e15.

Pezzotta, A. and Briscoe, J. (2022). Optimal control of gene regulatory networks for morphogen-driven tissue patterning. bioRxiv.

Pick, L., Schier, A., Affolter, M., Schmidt-Glenewinkel, T., and Gehring, W. J. (1990). Analysis of the ftz upstream element: Germ layer-specific enhancers are independently autoregulated. Genes and Development, 4(7):1224– 1239.

Reimer, A., Alamos, S., Westrum, C., Turner, M. A., Talledo, P., Zhao, J., and Garcia, H. G. (2021). Minimal synthetic enhancers reveal control of the probability of transcriptional engagement and its timing by a morphogen gradient. bioRxiv.

Sáez, M., Blassberg, R., Camacho-Aguilar, E., Siggia, E. D., Rand, D. A., and Briscoe, J. (2022). Statistically derived geometrical landscapes capture principles of decision-making dynamics during cell fate transitions. Cell Systems, 13(1):12–28.

Schier, A. F. and Gehring, W. J. (1992). Direct homeodomain–DNA interaction in the autoregulation of the fushi tarazu gene. Nature, 356(6372):804–807.

Schier, A. F. and Gehring, W. J. (1993). Analysis of a fushi tarazu autoregulatory element: Multiple sequence elements contribute to enhancer activity. EMBO Journal, 12(3):1111–1119.

Schroeder, M. D., Greer, C., and Gaul, U. (2011). How to make stripes: Deciphering the transition from nonperiodic to periodic patterns in Drosophila segmentation. Development, 138(14):3067–3078.

Soldatov, R., Kaucka, M., Kastriti, M. E., Petersen, J., Chontorotzea, T., Englmaier, L., Akkuratova, N., Yang, Y., Häring, M., Dyachuk, V., Bock, C., Farlik, M., Piacentino, M. L., Boismoreau, F., Hilscher, M. M., Yokota, C., Qian, X., Nilsson, M., Bronner, M. E., Croci, L., Hsiao, W.-Y., Guertin, D. A., Brunet, J.-F., Consalez, G. G., Ernfors, P., Fried, K., Kharchenko, P. V., and Adameyko, I. (2019). Spatiotemporal structure of cell fate decisions in murine neural crest. Science, 364(6444).

Soluri, I. V., Zumerling, L. M., Payan Parra, O. A., Clark, E. G., and Blythe, S. A. (2020). Zygotic pioneer factor activity of Odd-paired/Zic is necessary for late function of the Drosophila segmentation network. eLife, 9.

Sootla, A., Oyarzún, D., Angeli, D., and Stan, G.-B. (2016). Shaping pulses to control bistable systems: Analysis, computation and counterexamples. Automatica, 63:254–264.

Sprinzak, D., Lakhanpal, A., LeBon, L., Santat, L. A., Fontes, M. E., Anderson, G. A., Garcia-Ojalvo, J., and Elowitz, M. B. (2010). Cis-interactions between Notch and Delta generate mutually exclusive signalling states. Nature, 465:86–90.

Srinivasan, S., Hu, J. S., Currle, D. S., Fung, E. S., Hayes, W. B., Lander, A. D., and Monuki, E. S. (2014). A BMP-FGF morphogen toggle switch drives the ultrasensitive expression of multiple genes in the developing forebrain. PLoS Computational Biology, 10(2):e1003463.

Tkačik, G. and Gregor, T. (2021). The many bits of positional information. Development, 148(2):dev176065.

Verd, B., Crombach, A., and Jaeger, J. (2017). Dynamic maternal gradients control timing and shift-rates for Drosophila gap gene expression. PLoS Computational Biology, 13:e1005285.

Verd, B., Monk, N. A., and Jaeger, J. (2019). Modularity, criticality, and evolvability of a developmental gene regulatory network. eLife, 8:e42832.

Villaverde, A. F., Henriques, D., Smallbone, K., Bongard, S., Schmid, J., Cicin-Sain, D., Crombach, A., Saez-Rodriguez, J., Mauch, K., Balsa-Canto, E., Mendes, P., Jaeger, J., and Banga, J. R. (2015). BioPreDyn-bench: a suite of benchmark problems for dynamic modelling in systems biology. BMC Systems Biology, 9(1).

von Dassow, G., Meir, E., Munro, E. M., and Odell, G. M. (2000). The segment polarity network is a robust developmental module. Nature, 406(6792):188–192.

Waddington, C. (1957). The Strategy of the Genes. Routledge.

Wakimoto, B. T., Turner, F. R., and Kaufman, T. C. (1984). Defects in embryogenesis in mutants associated with the antennapedia gene complex of Drosophila melanogaster. Developmental biology, 102(1):147–172.

Wang, J., Zhang, K., Xu, L., and Wang, E. (2011). Quantifying the Waddington landscape and biological paths for development and differentiation. Proceedings of the National Academy of Sciences, 108(20):8257–8262.

Weiner, A. J., Scott, M. P., and Kaufman, T. C. (1984). A molecular analysis of fushi tarazu, a gene in Drosophila melanogaster that encodes a product affecting embryonic segment number and cell fate. Cell, 37:843–851.

Wieland, F.-G., Hauber, A. L., Rosenblatt, M., Tönsing, C., and Timmer, J. (2021). On structural and practical identifiability. Current Opinion in Systems Biology, 25:60–69.

Xiong, W. and Ferrell, J. E. (2007). A positive-feedback-based bistable ‘memory module’ that governs a cell fate decision. Nature, 448(7157):1076–1076.

Zernicka-Goetz, M., Morris, S. A., and Bruce, A. W. (2009). Making a firm decision: multifaceted regulation of cell fate in the early mouse embryo. Nature Reviews Genetics, 10(7):467–477.

Zhang, L., Radtke, K., Zheng, L., Cai, A. Q., Schilling, T. F., and Nie, Q. (2012). Noise drives sharpening of gene expression boundaries in the zebrafish hindbrain. Molecular Systems Biology, 8(1):613.

